# Gut microbial genomes with paired isolates from China signify probiotic and cardiometabolic effects

**DOI:** 10.1101/2023.09.25.559444

**Authors:** Pan Huang, Quanbing Dong, Yifeng Wang, Yunfan Tian, Shunhe Wang, Chengcheng Zhang, Leilei Yu, Fengwei Tian, Xiaoxiang Gao, Hang Guo, Shanrong Yi, Mingyang Li, Yang Liu, Qingsong Zhang, Wenwei Lu, Gang Wang, Bo Yang, Shumao Cui, Dongxu Hua, Xiuchao Wang, Yuwen Jiao, Lu Liu, Qiufeng Deng, Beining Ma, Tingting Wu, Huayiyang Zou, CGMR Consortium, Jing Shi, Haifeng Zhang, Daming Fan, Yanhui Sheng, Jianxin Zhao, Liming Tang, Hao Zhang, Wei Sun, Wei Chen, Xiangqing Kong, Lianmin Chen, Qixiao Zhai

## Abstract

The gut microbiome displays significant genetic differences between populations while systematic characterization of the genomic landscape of the gut microbiome in Asia populations remains limited. Here, we present the Chinese gut microbial reference (CGMR) set, comprising 101,060 high quality metagenomic assembled genomes (MAGs) of 3,707 non-redundant species paired with 1,376 live isolates from a national wide collection of 3,234 fecal samples across China. This improved reference set contains 987 novel species compared with existing resources worldwide. By associating MAGs with geographic and phenotypic characteristics, we observed regional-specific coexisting MAGs and MAGs with probiotic and cardiometabolic functionalities. We further conducted mice experiments to confirm the probiotic effects of two *Faecalibacterium intestinalis* isolates in alleviating constipation, the cardiometabolic influences of three *Bacteroides fragilis_A* isolates in obesity, and the functional potential of isolates from the two new species belonging to the genera *Parabacteroides* and *Lactobacillus* in host lipids metabolism. Our study not only expands the current microbial genomes with paired isolates but also demonstrates their probiotic and cardiometabolic effects on hosts, contributing to the mechanistic understanding of host-microbe interactions and the translation of microbiome-based personalized therapies.

## INTRODUCTION

Variations in gut microbial composition have been associated with human health and various diseases, including cardiovascular disease (CVD), obesity, inflammatory bowel disease (IBD), and diabetes ^1-3^. However, the functional potential of gut microbes underlying these associations is largely unexplored ^4^, as a substantial number of microbes have not yet been isolated, and their genomes are not well-profiled ^5-7^. Therefore, characterizing diverse microbial genomes and cultivating live microbes is an important step in decoding the role of microbes in human health and disease.

Recently, numerous efforts have been devoted to generating comprehensive microbial reference genomes by using large-scale metagenomic assembly of samples from diverse populations ^5–9^. Meanwhile, thousands of gut microbial isolates have also been cultivated, which contributes to the mechanistic understanding of host-microbe interactions and the probiotic potential in clinical translation ^10–13^. However, most of the metagenomic samples utilized in the above-mentioned studies are from European and American ancestries, and only around 10% are from Asia ^5, 6^. Microbial genomes reported from China are also limited in both sample size and specific region ^14^. Notably, prominent species are found in people worldwide but display substantial genetic differences between human populations ^5, 6^. Species exhibiting the most significant codiversification have independently developed traits associated with host dependence, emphasizing the need to comprehend the potential impact of population-specific microbial strains on microbiome-related disease phenotypes ^15^. Thus, expanding microbial genomes and isolates on a nationwide scale in China may act as a valuable addition to the field and have implications for the generalizability of microbiota-based therapeutics for the population.

Here, we present a set of 101,060 metagenomic assembled genomes (MAGs) from 3,234 randomly collected fecal samples from 30 provinces in mainland China, referred to as the Chinese gut microbial reference (CGMR) set. These MAGs can be clustered into 3,707 non-redundant species, of which 987 are newly reported species compared to existing databases, including CIBIO ^5^, UHGG ^6^, WIS ^7^, ELGG ^16^, IMGG ^14^ and GTDB ^17^. Additionally, we cultured a total of 1,376 MAG-paired live isolates, which included two isolates from two newly identified species. By associating MAGs with geographic and phenotypic characteristics, we observed regional-specific coexisting MAGs and MAGs with probiotic and cardiometabolic functionalities. We further conducted mice experiments to confirm the probiotic effects of two *Faecalibacterium intestinalis* isolates in alleviating constipation, the cardiometabolic influences of three *Bacteroides fragilis_A* isolates in obesity, and the functional potential of isolates from the two new species belonging to the genera *Parabacteroides* and *Lactobacillus* in host lipids metabolism. Our study not only expands the current microbial genomes with paired isolates but also demonstrates their probiotic and cardiometabolic effects on hosts, contributing to the mechanistic understanding of host-microbe interactions and the translation of microbiome-based personalized therapies.

## RESULTS

### The genomic landscape of gut microbiome in Chinese

To comprehensively characterize the genomic landscape of gut microbes in Chinese on a nationwide scale, we collected fecal samples from 3,234 participants across 30 provinces in mainland China (**Figure 1A**). Through metagenomic sequencing, we obtained an average of 65 million paired reads (20 GB) for each sample. We utilized a single sample metagenomic de novo assembly pipeline (**Figure 1B**), building upon existing workflows ^5, 6^. Following our pipeline (**Figure 1B**), a total of 292,550 metagenomic assembled genomes (MAGs) were obtained, out of which 101,060 met the minimum information about a metagenome-assembled genome (MIMAG) standard ^18^. These MAGs exhibited a completeness of ≥ 50% and a contamination rate of < 5% (completeness - 5 × contamination, QS > 50). It is worth noting that the assembled genomes were of high quality, with a median completeness of 88.28% and a contamination rate of 0.98% (**Figure 1C**, **Table S1**). The median size of the 101,060 MAGs was 2.20 MB (interquartile range, IQR = 1.75-2.71 Mb), with N50 values ranging from 1.8 kilobases to 1.05 Mb (**Table S1**). We refer to this collection as the Chinese Gut Microbial Reference (CGMR) set.

**Figure 1.**
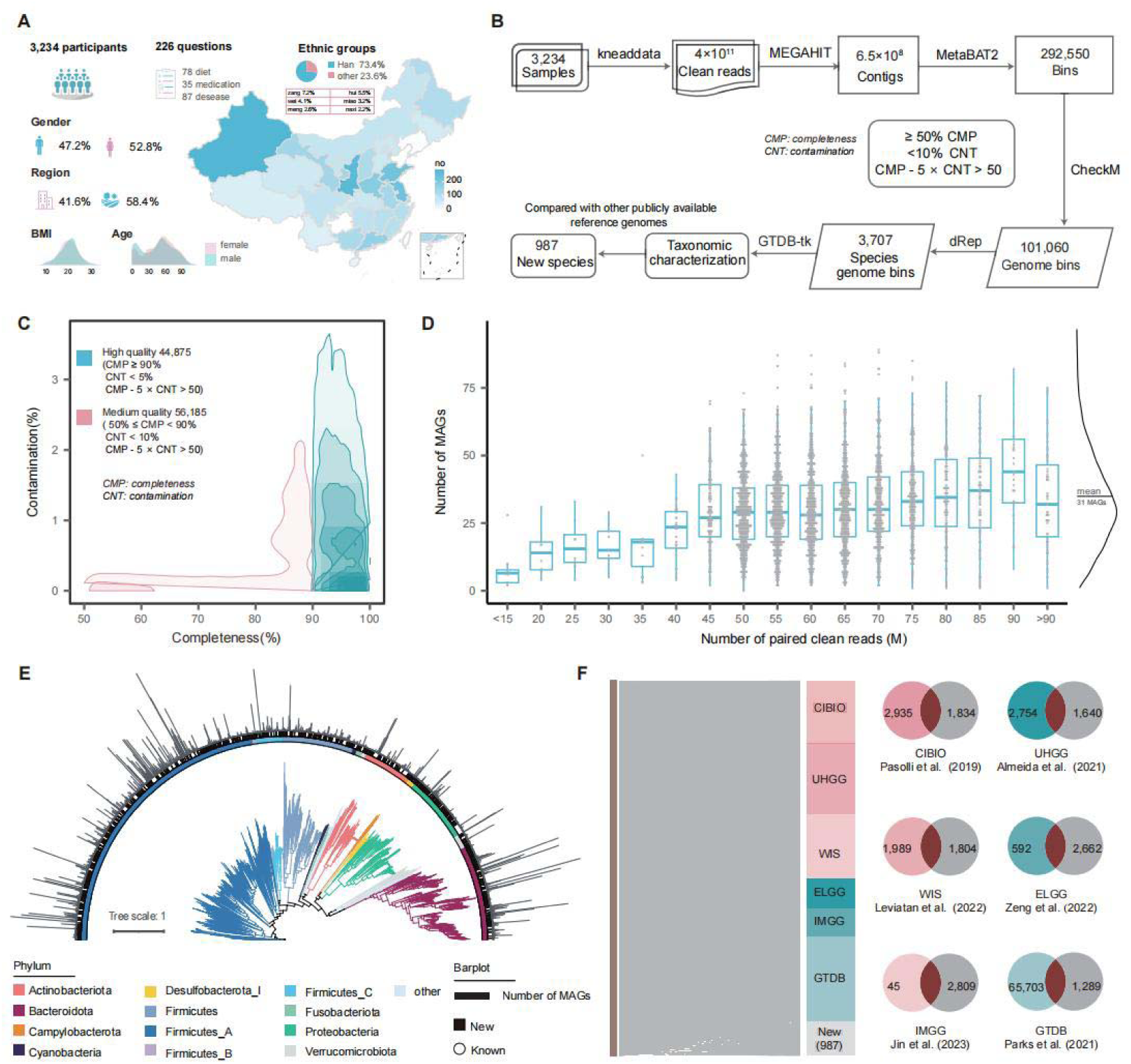
The Chinese gut microbiome reference set (CGMR). **A.** Geographic distribution and phenotypic characteristics of the 3,234 samples retrieved from mainland China. The shades of blue represent the number of samples collected per province. Histograms depict the distribution of age and BMI of the participants. **B.** Overview of the metagenomic assembly strategy and datasets used for the reconstruction and comparison of microbial genomes. In total, 101,060 metagenomic assembled genomes (MAGs) were generated, representing 3,707 species, with 987 species newly reported compared to existing datasets. **C.** Genomic quality distribution of the 101,060 MAGs. Completeness and contamination values estimated by CheckM are shown for medium-quality (pink) and high-quality (blue) genomes. The intensity of the colors indicates the density of MAGs. **D.** Total number of MAGs obtained per sample at different sequencing depths. Each dot represents one sample, with the x-axis indicating the number of paired clean reads and the y-axis indicating the number of MAGs obtained per sample. Box plots display medians, the first and third quartiles of the MAG numbers. The upper and lower whiskers extend no further than 1.5×IQR (interquartile range), respectively. Outliers are plotted individually. **E.** Phylogenetic tree of all 3,707 representative species-level MAGs. The colors on the branches represent the bacterial phyla, while the color on the outer circle indicates newly identified species compared to existing genomic references. The length of the outer bars represents the total number of MAGs obtained for each representative species. **F.** Comparison of CGMR to 6 existing genome collections. The sankey plot illustrates the links between CGMR MAGs and the existing microbial genome sets. Venn diagrams show the unique MAGs of CGMR when compared with others, with the respective numbers indicated.

Of note, our sequencing data has approximately four times higher read depth compared to recent studies ^5–7^. On average, we obtained 31 MAGs per sample, which is twice the number obtained in previous studies ^5–7^. We also observed a positive correlation between read depth and the number of MAGs obtained per sample (**Figure 1D**). It is noteworthy that the total number of MAGs obtained per sample reached an optimal level at around 80 million reads (**Figure 1D**), indicating that deep sequencing was crucial to obtain an adequate quantity of MAGs.

Next, we analyzed the taxonomy of these MAGs by clustering them using an average nucleotide identity (ANI) threshold of 95%, resulting in 3,707 representative prokaryotic species (**Figure 1E**, **Table S2**). With GTDB-Tk ^19^, taxonomic annotatable genomes belong to 28 phyla, 37 classes, 93 orders, 215 families, and 972 genera (**Table S2**). Dominant phyla include *Firmicutes_A*, *Bacteroidota*, *Proteobacteria*, and *Actinobacteriota* (**Figure 1E**). Our CGMR set has fewer bacterial genera and families compared to WIS (2365 genra and 627 family) and CIBIO (2640 genra and 778 family), although the number of representative species is identical. This suggests the potential uniqueness of the gut microbial genomic landscape in the Chinese population.

### CGMR set expands the current microbial genomes by adding 987 new species

Microbial genomes are essential resources for exploring microbial diversity and understanding microbial community structure and function. However, the representation of samples from the Chinese population has been limited in previous studies despite the reporting of over 200,000 metagenomic assembled genomes (MAGs) from populations worldwide ^5–7, 14^.

To demonstrate the added value of our CGMR set, we compared the 3,707 species-level representative genomes with existing microbial genomes obtained from studies including CIBIO ^5^, UHGG ^6^, WIS ^7^, ELGG ^16^ and IMGG ^14^. Using a 95% ANI threshold, we found that 1,834, 1,640, 1,989, 2,662, and 2,809 of the CGMR set genomes were novel compared to CIBIO, UHGG, WIS, ELGG, and IMGG, respectively (**Figure 1F**). Importantly, out of the 3,707 species-level representative genomes, 1,371 (37%) were not covered in the above-mentioned studies (**Figure 1F**, **Table S3**). This highlights the uniqueness of the gut microbiome in the Chinese population.

Furthermore, we mapped the 1,371 unique species-level representative MAGs using GTDB-Tk ^17^. It was found that 987 of these MAGs could not be taxonomically classified at the species level, 110 could not be annotated at genus level, and one of them (FNXYCHL24.61) could not even be assigned to existing order levels. This indicates that our CGMR set expands the current microbial genomes by adding at least 987 potential new species. These genomes serve as critical resources for exploring microbial diversity and understanding microbial community structure and function in the Chinese population.

### Geographic characteristics of CGMR and keystone species

China exhibits significant geographic diversity, with various landscapes across different regions. The eastern plains and southern coasts are characterized by fertile lowlands and foothills, while the southern areas feature hilly and mountainous terrain. The western and northern parts of the country consist of sunken basins, rolling plateaus, and towering massifs. These geographic features give rise to diverse regional-specific environments and lifestyles that can potentially influence the distribution of the gut microbiome.

To investigate the impact of geography on the gut microbiome, we divided the country into 7 regions based on their geographic locations (**Figure 2A**). By further analyzing the species-level representative MAGs, we identified 115 MAGs that had a prevalence rate of more than 10% in at least one region (**Table S4**). Notably, we found 45 MAGs (from 34 genera) that were commonly present in all 7 regions (**Figure 2A**). Among these common species, several have been suggested to possess probiotic potential, such as *Akkermansia muciniphila*, *Faecalibacillus intestinalis*, *Bifidobacterium longum* and *Bifidobacterium pseudocatenulatum* (**Table S4**). These findings highlight the presence of beneficial microbes that may contribute to the health and balance of the gut microbiome across different regions of China.

**Figure 2.**
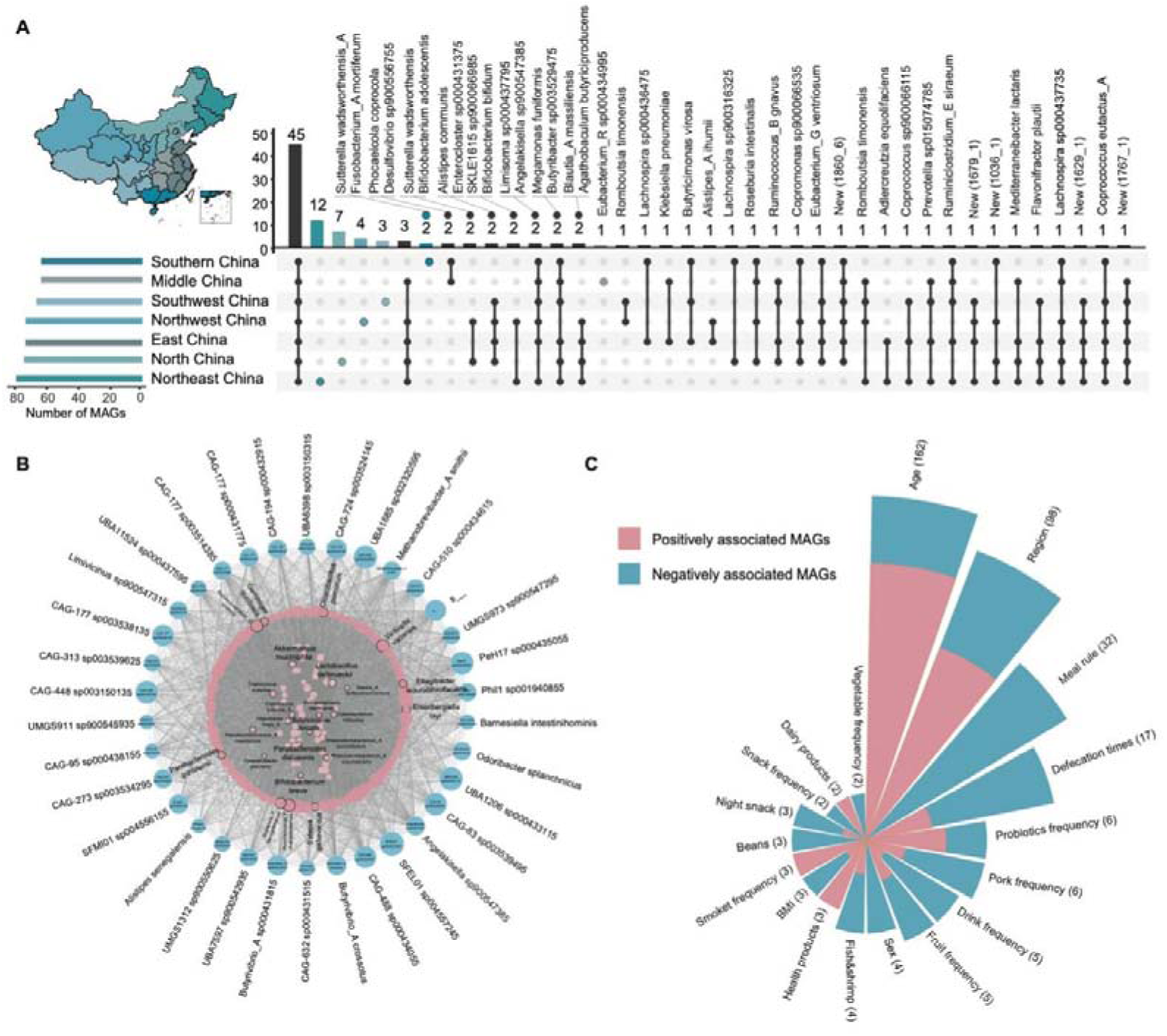
Regional-specific and keystone MAGs and associations with host phenotypes. **A.** Geographic distribution of prevalent species-level MAGs. The map shows the seven regions in China, with colors indicating the prevalence of species-level MAGs. The left panel displays horizontal bars indicating the total number of prevalent species-level MAGs in each region. The vertical bars represent the number of prevalent species-level MAGs shared between specific regions highlighted with colored dots in the lower panel. Different shades of blue indicate prevalent species-level MAGs specific to only one region. **B.** Co-occurrence network based on the prevalence of species-level mags. Each dot represents one species, with its name marked, and gray lines connect two dots to indicate significant species-level co-occurrences (FDR < 0.05). The top 1% keystone species are highlighted in blue, while the top 5% are in pink. **C.** Phenotypic associations with the prevalence of species-level MAGs. Bars represent host phenotypes, sorted by the number of significant associations. The association directions are color-coded, with pink indicating a positive association and blue indicating a negative association. The length of each bar indicates the number of significant associations at FDR < 0.05.

Notably, we identified 27 species-specific MAGs, with 19 of them being specific to the northeast and north regions of China (**Figure 2A**). Despite having the fewest samples (**Figure S1**), the northeast of China exhibited the highest number of regional-specific species, including 12 species from the genera *Anaerobutyricum*, *Blautia*, *Butyribacter*, *Clostridium*, *Coprococcus*_A, *Eggerthella*, *Evtepia*, *Lachnospira*, *Mediterraneibacter*, *Methanobrevibacter*_A, *Ruminococcus*_E, *Turicibacter* (**Figure 2A**, **Table S4**). Noteworthy, half of these species belong to the *Lachnospiraceae* family. *Lachnospiraceae* is a well-documented intestinal symbiotic bacteria known to produce beneficial metabolites for the host, such as short-chain fatty acids (SCFAs) ^20^. The unique geographic position of northeastern China (118°E-135°E, 48°N-55°N) contributes to its temperate continental climate, which is influenced by cold air from Siberia. This region experiences long and cold winters, along with short and rainy summers. Interestingly, studies have suggested that the abundance of *Lachnospiraceae* can be influenced by temperature and/or wind speed ^21, 22^. This may potentially explain the prevalence of *Lachnospiraceae* in the population of northeastern China, highlighting the influence of climatic factors on the composition of the gut microbiome in this region.

In addition to the region-specific species MAGs, we also investigated microbial taxa that play a potentially critical role in the organization and maintenance of the gut ecosystem. These taxa are characterized by their central position in microbial co-occurrence networks ^1, 23^ and are referred to as keystone species. We identified a total of 95,213 significant co-occurrences involving 3,462 species-level MAGs (FDR < 0.05, **Figure 2B**, **Table S5**). Among these co-occurrences, we defined 348 MAGs with high connectivity (top 10%) as keystone species, representing 210 genera (**Table S5**). These keystone species are likely crucial for maintaining the stability of the gut ecosystem and understanding the significance of the gut microbiome in human health and disease. Notably, certain keystone species, such as *Butyrivibrio*, have been associated with multiple cardiometabolic diseases in recent studies ^1, 24^. Collectively, these findings highlight the unique geographical and ecological characteristics of the gut microbiome in the Chinese population, emphasizing the importance of considering both regional-specific species and keystone taxa in understanding the complexities of the gut ecosystem and its impact on human health.

### Prevalence of MAGs associated with multiple host phenotypes

In addition to the geographical characteristics, we collected information on 210 phenotypes from the study participants. These phenotypes encompassed anthropometric traits (e.g., age, sex, and body mass index BMI), residential location (urban and rural), dietary habits, lifestyles, as well as disease history and medication use (**Table S6**). The continuous traits, such as BMI, followed a normal distribution (**Figure 1A**). For binary traits, such as residential locations, an equal distribution was observed among the seven regions (**Figure 2A**), with 41.6% residing in urban areas and 58.4% in rural areas (**Figure 1A**; **Table S6**).

By analyzing the associations between the prevalence of MAGs and various phenotypic traits, we identified a total of 371 significant associations (FDR < 0.05, **Figure 2C**; **Table S7**). Age showed the highest number of associations, with 162 MAGs from 114 genera being correlated with age. Notably, several of these MAGs belonged to probiotic species, such as *Bifidobacterium breve*, *Bifidobacterium longum*, *Bifidobacterium bifidum*, *Bifidobacterium pseudocatenulatum*, and *Bifidobacterium animalis*. These findings are consistent with previous population studies that have suggested a negative association between the abundance of *B. bifidum* and *B. longum* and age ^25^.

Additionally, we identified 98 MAGs associated with residential location (urban or rural), primarily enriched for species from 77 genera (FDR < 0.05, **Table S7**). Specifically, 32 MAGs were more prevalent in urban areas, while 66 MAGs were more prevalent in rural areas. MAGs associated with rural areas were enriched for species from the genera *Prevotella*, *Eubacterium*, *Phascolarctobacterium Coprococcus*, *Agathobacter*, *Alistipes* and *Cryptobacteroides*, as well as taxa related to fiber, amino acid, and vitamin metabolism. In contrast, industrialized urban populations were characterized by a dominance of *Bifidobacterium*, *Streptococcus*, *Staphylococcus*, *Clostridium*, *Enterococcus* and other members of *Firmicutes* (**Table S7**). Additionally, MAGs belonging to *Staphylococcus aureus*, *Clostridium paraputrificum*, and *Enterococcus faecalis* were only found in urban populations, suggesting potential health risks associated with urbanization ^26–28^.

Furthermore, we observed 16 MAGs linked to defecating frequency and 3 MAGs associated with BMI. For example, the prevalence of *F. intestinalis* was positively associated with defecating frequency (r = 0.10; P = 3.17×10^-5^), while *Bacteroides fragilis_A* showed an association with BMI (r = −0.13; P = 1.39×10^-5^). Regarding disease and medication associations, we observed limited associations due to the small number of participants with such records (**Table S6**), which resulted in reduced statistical power. Overall, these findings demonstrate significant associations between the prevalence of MAGs and various host phenotypes, including age, residential location, defecating frequency, and BMI. Notably, probiotic species were identified among the age-related associations, emphasizing their potential role in age-related gut microbiome dynamics.

### A set of 1,376 MAG-paired bacterial isolates enabling functional exploration

The cultivation of live microbial isolates allows for a more comprehensive investigation of the gut microbiota, including the identification of metabolic functionalities associated with diseases and the characterization of next-generation probiotics. To achieve this, we isolated live microbes from 982 fecal samples, 664 of which were paired with fecal metagenomic sequencing data. With 27 different culture media (**Figure S2**), we successfully obtained a total of 1,376 isolates. Subsequently, we performed whole-genome sequencing on these isolates, resulting in high-quality genomes with an average completeness 99.4% and contamination 1.1% (**Figure 3A**; **Table S8**).

**Figure 3.**
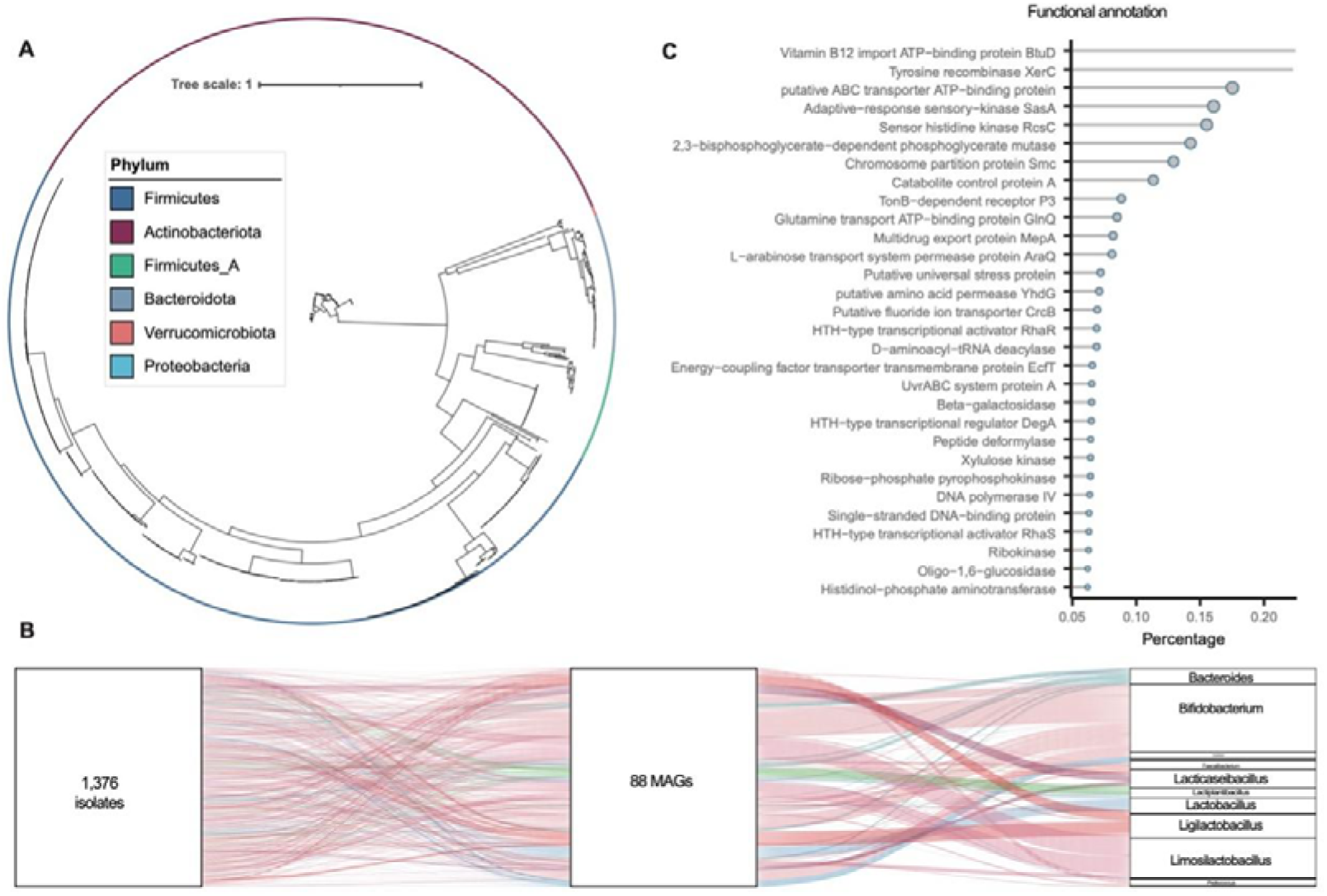
Overview of the 1,376 bacterial isolates. **A.** Phylogenetic tree of all the 1,376 live isolates based on their draft genomes. The phylogenetic tree depicts the relationships among the 1,376 bacterial isolates, with the colors on the outer circle representing the phyla to which each isolate belongs compared to existing genomic references. **B.** Isolate-MAG Pairs. The parallel coordinates chart illustrates the connections among the 1,376 isolates (left panel), 88 species-level representative MAGs (middle panel), and their corresponding genera information (right panel). The curved lines that connect the panels indicate the linkages, with colors corresponding to different microbial genera. **C.** Functional annotation of genes in the draft genome of isolates. The y-axis indicates the encoded functions of annotatable genes, while the x-axis indicates the percentage of genes that encode a certain function.

We further conducted a comparison of these genomes with our CGMR set and found that all of them exhibited a match to CGMR MAGs at a 95% ANI threshold (**Table S9**). This included 69 species from 6 phyla and 26 genera (**Figure 3B**, **Table S9**), as well as 2 isolates representing newly characterized species in our study. Notably, many of these isolates belonged to probiotic species such as *Bifidobacterium*, *Lactobacillus*, and *Lacticaseibacillus* species, as well as potential next-generation probiotics including *Bacteroides*, *Faecalibacterium*, and *Clostridium* species (**Figure 3B**, **Table S9**).

Using standard tools, we annotated the genomes of these isolates and identified a total number of 3,501,359 gene segments, which may not necessarily be annotated genes themselves. Among these, 12,700 genes were uniquely annotated, counting each gene only once even if it appeared in multiple species. Approximately 41% of the identified genes were hypothetical, while the annotated genes encompassed a wide range of functionalities including vitamin and multidrug transport, DNA topology modulation, as well as the biosynthesis of bile acids, amino acids, and SCFAs (**Figure 3C**, **Table S10**). These findings demonstrate how our collection expands the taxonomic diversity of cultivable microbes in the human gut, providing valuable live isolates for targeted mechanism validation.

### Probiotic effects of *Faecalibacterium intestinalis* in alleviating constipation

Constipation is a chronic bowel disorder that can be attributed to various factors including lifestyle, psychological, and behavioral factors ^29^. Previous studies have associated *Faecalibacterium* with faster colonic transit ^30^. Here, we observed significant associations between the prevalence of 17 MAGs and defecating frequency (FDR < 0.05, **Figure 2C**), and one of the strongest associations was found with *F. intestinalis* (r= 0.103, P= 3.17×10^-3^, **Figure 4A; Table S7**). To further validate this observation, we selected two isolates of *F. intestinalis* (F1 and F2) from our collection of 1,376 isolates to evaluate their efficacy in alleviating constipation induced by loperamide. We employed a specific pathogen-free (SPF) C57BL/6J mice model (**Figure S3**). As a positive control known for its beneficial effects on constipation, we also included *Bifidobacterium animalis ssp. lactis* BB-12 ^31^.

**Figure 4.**
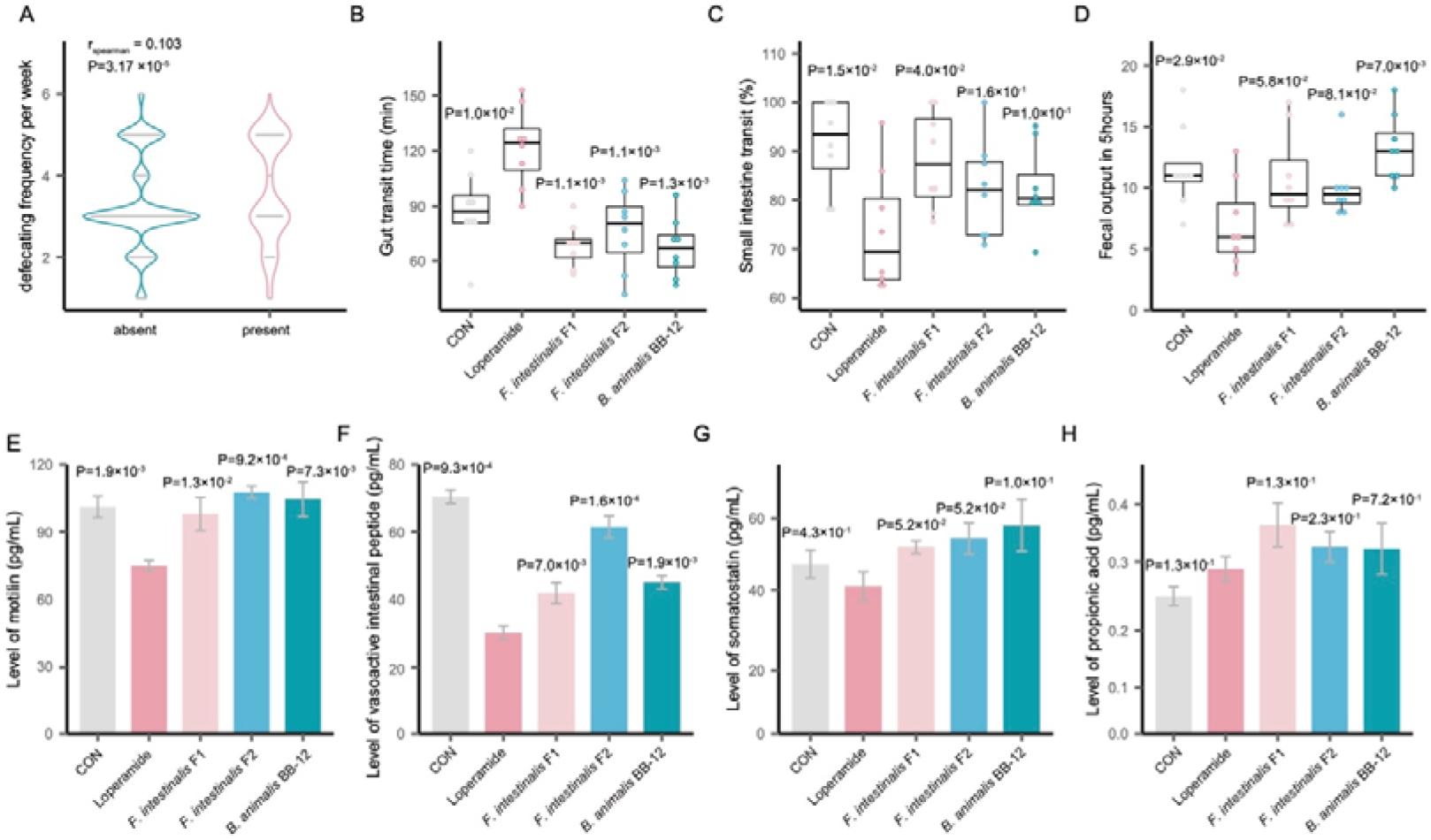
*Faecalibacterium intestinalis* isolates alleviate constipation. **A.** Defecation frequency difference between participants with and without *Faecalibacterium intestinalis* MAG. Each dot represents an individual, and the violin plot illustrates the distribution of defecation frequency per week. The Spearman test is used to assess the difference between the two groups. **B.** Differences in gut transit time between groups. **C.** Differences in small intestine transit between groups. **D.** Differences in 5-hour fecal output between groups. **E.** Differences in motilin levels between groups. **F.** Differences in vasoactive intestinal peptide levels between groups. **G.** Differences in somatostatin levels between groups. **H.** Differences in propionic acid levels between groups. For panels **B-D**, each dot represents one sample and is colored differently based on the group. Box plots display the medians and the first and third quartiles of the corresponding traits’ levels. The upper and lower whiskers extend to the largest and smallest values no further than 1.5 × IQR (interquartile range), respectively. Outliers are plotted individually. For panels **E-H**, bar plots represent the mean levels of the traits within groups, with standard deviations indicated above the bars. The difference between groups is assessed using the Wilcoxon test, and the corresponding P-value is provided.

Compared to the control group (CON), the mice treated with loperamide (LOP) exhibited typical symptoms of constipation. This was evident from the increased whole gut transit time (P = 0.010, **Figure 4B**), decreased small intestine transit (P = 0.015, **Figure 4C**), and reduced fecal output (P = 0.029, **Figure 4D**). Importantly, the intragastric administration of *F. intestinalis* isolates F1 and F2, as well as *B. animalis ssp. lactis* BB-12, resulted in a significant reduction in whole gut transit time compared to the LOP group (P < 0.05). Furthermore, the small intestine transit was improved with the administration of *F. intestinalis* F1 (**Figure 4C**).

Previous studies have demonstrated that motilin and vasoactive intestinal peptide (VIP) produced in the gut are beneficial for intestinal motility ^32, 33^. Additionally, SCFAs, such as propionic acid, acetic acid, and butyric acid, have been observed to be negatively associated with constipation ^34^. In our study, we examined the levels of these molecules and found that the intragastric administration of *F. intestinalis* isolates F1, F2, and *B. animalis ssp. lactis* BB-12 significantly increased the levels of motilin, VIP and SS (**Figure 4E-G**). Furthermore, the *F. intestinalis* F1 and F2 groups showed higher levels of propionic acid compared to the CON group (**Figure 4H**). These findings suggest that the use of *F. intestinalis* isolates as probiotics has the potential to alleviate constipation by influencing the levels of motilin, VIP, and SCFAs.

### BMI-related *Bacteroides fragilis_A* influences cardiometabolic traits in obesity

In addition to constipation, we also observed a negative association between the prevalence of *B. fragilis_A* and host BMI (**Figure 2C**, **Figure 5A**), indicating a potential beneficial role of *B. fragilis_A* in cardiometabolic health. We administered three *B. fragilis_A* isolates (B6, B7, and B8) from the 1,376 isolates to mice fed a high-fat diet (HFD), with *B. fragilis* ATCC25285 (BATCC, **Figure S4**), a promising next-generation probiotics ^35, 36^, serving as the reference group.

**Figure 5.**
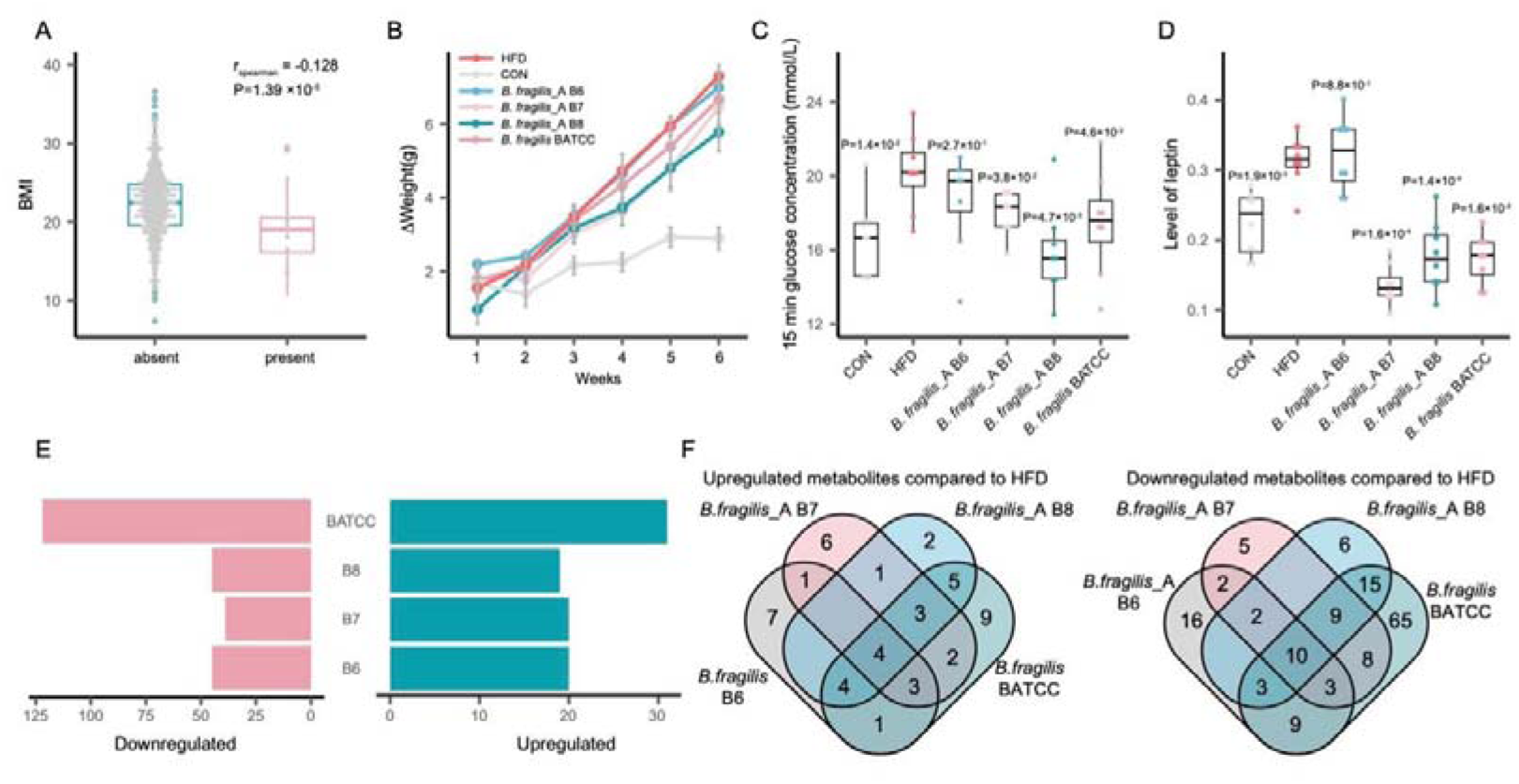
*Bacteroides fragilis*_A isolates influence cardiometabolic traits in obesity. **A.** Difference in BMI between participants with and without *Bacteroides fragilis*_A MAG. Each dot represents an individual, and the box plot with dots represents the distribution of BMI. The Spearman test is used to assess the difference between the two groups. **B.** Differences in changes of body weight between groups over 6 weeks. **C.** Differences in 15-minute OGTT (oral glucose tolerance test) glucose levels between groups. **D.** Differences in leptin levels between groups. For panels **A** and **C-D**, each dot represents one sample, colored differently based on the group. Box plots show medians and the first and third quartiles of the corresponding traits’ levels. The upper and lower whiskers extend to the largest and smallest values no further than 1.5 × IQR, respectively. Outliers are plotted individually. For panels **B-D**, the difference between groups is shown by the Wilcoxon test, and the corresponding P-value is provided. **E.** Number of differential metabolites compared with the HFD (high-fat diet) group. The x-axis indicates the number of differential metabolites at FDR<0.05 by the Wilcoxon test. **F.** Overlap of Differential Metabolites between Groups.

After 6 weeks of feeding, we observed that mice fed the HFD had significantly higher body weight from week 4 to 6 compared to the control group (CON) fed a chow diet (P < 1.1 × 10^-2^, **Figure 5B**, **Table S11**). When comparing the HFD group with the other groups fed different *B. fragilis* isolates, we did not observe significant differences in body weight (**Figure 5B**, **Table S11**). However, we found that glucose levels were lower in B7, B8, and BATCC at 15 minutes during the oral glucose tolerance test (OGTT) (**Figure 5C**, **Table S11**), and the levels of leptin were also lower in those groups (**Figure 5D**, **Table S11**).

To understand the functional roles of these isolates in influencing host cardiometabolic traits, we profiled the metabolome of feces in the colon and observed significant differences in 428 out of 1,437 metabolites between the CON and HFD groups (FDR < 0.05, **Table S12**). These differences included 194 upregulated and 234 downregulated metabolites in the HFD group (FDR < 0.05, **Table S12**). Notably, 20, 20, 19, and 31 HFD upregulated metabolites showed lower levels in B6, B7, B8, and BATCC, respectively, compared to HFD (**Figure 5E-F**, **Table S12**). Similarly, 45, 39, 45, and 122 HFD downregulated metabolites showed higher levels in B6, B7, B8, and BATCC, respectively, compared to HFD (**Figure 5E-F**, **Table S12**). The intervention of bacterial isolates resulted in alterations in 258 differential metabolites between the CON and HFD groups, including carbohydrate conjugates, amino acids, fatty acids, and secondary bile acids (**Table S12**).

We further investigated if the differential metabolites altered by the four isolates could be replicated *in vitro* using the gut microbiota medium (GMM) ^37^ for fermentation. Remarkably, 58 out of the 258 differential metabolites could be measured in the *in vitro* fermentation, and 41 of them also showed differences between the blank GMM and medium with the four isolates (P< 0.05, **Table S13**). Interestingly, out of the 41 metabolites, 10 exhibited differences only in B6, B7, or B8, but not in BATCC (**Table S13**). For instance, we observed that B8 was involved in the biosynthesis of gamma-glutamyl leucine, a dipeptide associated with decreased mortality ^38^. Animal studies suggest that gamma-glutamyl leucine is an indicator of anti-obesogenic metabolism ^39^, as obesity is a strong risk factor for cardiometabolic disorders. These findings highlight the potential of BMI-related *Bacteroides fragilis_A* as a modulator of cardiometabolic traits in obesity.

### Isolates from new species modulate lipids metabolism in germ-free mice

Among the 987 newly recovered species in our CGMR set, we obtained live isolates from two new species: *Parabacteroides* (FGD16K12) and *Lactobacillus* (FHNXY56M7), referred to as P.new and L.new, respectively. Although we did not observe any significant association between the prevalence of these two novel MAGs and host phenotypes, our aim was to explore their potential roles in host metabolic health. For this purpose, we conducted mono-colonization experiments using germ-free mice (**Figure S5**). After four days of gavage, fresh fecal samples were collected and plated at 1, 5, and 10 days after gavage to ensure the stable colonization of the two isolates in germ-free mouse models (**Figure 6A**). Blood and colon tissues were collected to measure metabolites and gene expression, respectively.

**Figure 6.**
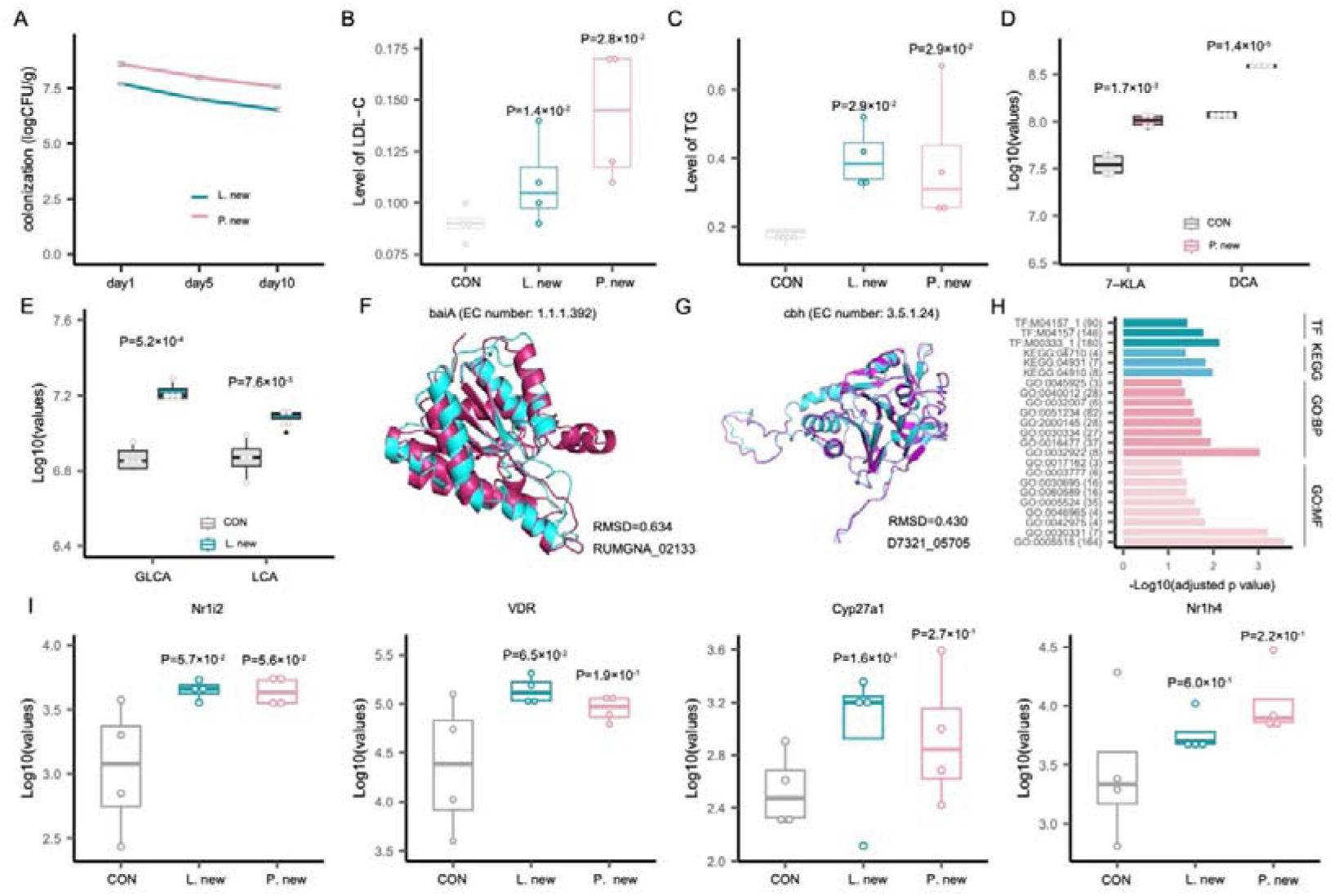
Isolates from new *Parabacteroides* and *Lactobacillus* species regulate lipids metabolism. **A.** Mono-colonization of isolates from new *Parabacteroides* and *Lactobacillus* species. Lines show the colonized number of bacteria at different days, with standard deviations marked in grey. **B.** Differences in LDL-C (low-density lipoprotein cholesterol) levels between groups. **C.** Differences in TG (triglyceride) levels between groups. **D.** *Parabacteroides* isolate-induced differences in bile acids. **E.** *Lactobacillus* isolate-induced differences in bile acids. For panels **B-E**, each dot represents one sample, colored differently based on the group. Box plots show medians and the first and third quartiles of the levels of the corresponding traits. The upper and lower whiskers extend to the largest and smallest values no further than 1.5 × IQR, respectively. Outliers are plotted individually. The difference between groups is shown by the Wilcoxon test, and the corresponding P-value is provided. **F.** Comparison of protein structures encoded by potential bile acid metabolism gene *baiA* from *Parabacteroides* FGD16K12. **G.** Comparison of protein structures encoded by potential bile acid metabolism gene *cbh* from *Lactobacillus* FHNXY56M7. For panels **F-G**, protein structure prediction is based on AlphaFold2. The resulting protein structures (blue) are then compared with confirmed bile acid metabolism proteins (purple) using PyMOL. The root-mean-square deviation (RMSD) value was calculated to evaluate the structural similarity between the modeled proteins and the confirmed proteins. **H.** Functional enrichment of differential genes in colon tissue of the L.new group. The y-axis indicates the enriched functions of differential genes, while the x-axis indicates the adjusted P-values.

For the serum metabolome, we identified 41 and 155 differential metabolites in P.new and L.new, respectively (**Table S14**). Remarkably, 11 and 69 of these metabolites were classified as lipids (**Table S14**) based on the annotation from HMDB ^40^ and LipidMaps ^41^. These findings partially align with the observed differences in low-density lipoprotein cholesterol (LDL-C) and total triglyceride (TG) levels between the two groups, respectively (**Figure 6B-C**).

Gut microbes have the potential to influence host lipid metabolism through various molecules, such as secondary bile acids ^42^. In our study, we observed higher levels of deoxycholic acid (DCA) and 7-ketolithocholic acid (7-KLA) in the P.new group, while lithocholic acid (LCA) and glycolithocholic acid (GLCA) were higher in the L.new group, compared to the control group (FDR<0.05, **Figure 6D-E**, **Table S14**). To investigate the capacity of the new isolates in bile acid metabolism, we analyzed their genomes and found that both isolates contain genes related to secondary bile acid synthesis. Specifically, *Parabacteroides* FGD16K12 harbors the *baiA* gene (EC number: 1.1.1.392) (**Figure 6F**), while *Lactobacillus* FHNXY56M7 has the *cbh_1* (EC number: 3.5.1.24) and *cbh_2* (EC number: 3.5.1.24) genes (**Figure 6G**).

Bile acids are known to regulate lipid metabolism through the activation of nuclear receptors, including farnesoid X receptor (FXR), vitamin D receptor (VDR), and pregnane X receptor (PXR) ^43^. At the transcriptional level, we observed differential expression of 253 and 532 genes in the colon tissues of the P.new and L.new groups, respectively (P< 0.01, Fold change> 2, **Table S15**). To gain further insight, we conducted gene functional enrichment analysis on the differentially expressed genes. In the L.new group, we identified a total of 28 enriched pathways, several of which are related to bile acid metabolism, such as nuclear estrogen receptor binding (GO:0030331) and nuclear retinoid X receptor binding (GO:0046965) (**Table S16**, **Figure 6H**). Estrogen regulates the expression of *CYP7B1* through insulin/phospho-inositol-3 kinase signaling ^44^, and the retinoid X receptor (RXR), as part of the RAR/RXR heterodimer, regulates the expression of the sodium taurocholate cotransporting polypeptide (NTCP). RXR inhibits NTCP while simultaneously inducing the efflux of bile acids into bile by upregulating the farnesoid X receptor (FXR) ^45, 46^. We didn’t observe much significant functional enrichment of in the P.new group.

We then analyzed the expression of genes involved in bile acid metabolism, including key genes in the classical synthesis pathway (*Cyp27a1*), as well as the bile acid receptors *FXR*, *PXR*, and *VDR*. Interestingly, all of these genes were found to be upregulated in both groups treated with the new isolates (**Figure 6I**). Furthermore, we also examined other potential bile acid genes encoding enzymes, transporters, and transcription factors, and significant differences were observed (**Figure S6**, **Table S15**). Overall, our data provide evidence that the isolates from these new species may modulate lipid metabolism through secondary bile acids.

## DISCUSSION

Prominent microbial species are found worldwide in the human gut but exhibit significant genetic variations across populations ^5, 6^, emphasizing the need to understand the potential role of population-specific microbial strains in microbiome-related disease phenotypes ^15^. However, the systematic characterization of the genetic landscape of the gut microbiome in the Chinese population remains limited. In this study, we present a comprehensive collection of 101,060 high-quality metagenomic assembled genomes (MAGs) representing 3,707 non-redundant species from a nationwide collection of 3,234 fecal samples in China. This collection, referred to as the Chinese gut microbial reference (CGMR) set, allows us to characterize the geographical specificity and ecological characteristics of the gut microbiome in the Chinese population, highlighting the importance of considering microbial strains in comprehending the complexities of the gut ecosystem and its impact on human health. Additionally, we employed 27 culture media to isolate 1,376 live strains and paired their MAGs with whole genomic sequencing data. This valuable resource enables us to functionally validate the observed associations between MAGs and phenotypes, including the probiotic and cardiometabolic effects of specific microbial strains.

Large-scale construction of microbial genomes has primarily focused on European and American samples ^6^. To address this gap and expand the current microbial reference genomes, we compared our dataset with recently reported large datasets of microbial genomes, including CIBIO ^5^, UHGG ^6^, WIS ^7^, ELGG ^16^, IMGG ^14^ and GTDB ^17^. Through this comparison, we have identified and added 987 new species-level representative metagenomic assembled genomes (MAGs) to the existing reference genomes. This significant expansion contributes to a more comprehensive and diverse representation of microbial genomes, particularly in non-European and non-American populations.

To identify the key microbial species in the Chinese population, we constructed a gut microbial co-occurrence network using the presence/absence information of species-level representative MAGs. Through our analysis, we discovered the presence of distinct keystone species, including potential probiotic species like *Akkermansia muciniphila*, *Faecalibacterium intestinalis*, *Bifidobacterium longum*, and *Bifidobacterium pseudocatenulatum*. Notably, these species are not commonly found in European individuals ^1^. This highlights the unique composition of the gut microbiome in the Chinese population and suggests the presence of potentially beneficial microbes that could contribute to gut health and overall well-being.

By conducting an analysis that associates MAGs with geographic and phenotypic characteristics, we discovered regional-specific MAGs, indicating significant variations in the composition of the gut microbiota across different regions in China. Notably, apart from East China, each region exhibited its distinct set of MAGs, which can be attributed, at least in part, to the developed economy and high population mobility in East China. Particularly interesting is the observation that Northeast China harbors the highest number of unique species, which may be influenced by the region’s specific climatic environment ^21, 47^. These findings highlight the impact of geographical factors on shaping the diversity and composition of the gut microbiota in different regions of China.

Furthermore, our study encompassed participants from various age groups, with the highest proportion being in the 50-60 age range. Notably, we included a significant number of infants (less than 3 years old, 10%) and elderly individuals over 84 years old (5.8%) in the study population. Correlation analysis revealed 162 MAGs that potentially exhibit associations with age. Additionally, we observed the presence of MAGs associated with 29 phenotypes, including defecation frequency and BMI. These findings suggest that specific microbial strains may play a role in modulating healthy aging and disease development.

To validate the associations between MAGs and phenotypes, we employed 34 different culture media for bacterial isolation and successfully obtained a total of 1,376 isolates. Among these isolates were two from previously uncharacterized species belonging to the genera *Parabacteroides* and *Lactobacillus*. Utilizing mouse models, we confirmed the probiotic effects of *Faecalibacterium intestinalis* isolates in alleviating constipation, the cardiometabolic influences of *Bacteroides fragilis* isolates in obesity, and the functional potential of the newly discovered *Parabacteroides* and *Lactobacillus* species isolates in host lipid metabolism via bile acids.

Taken together, our study not only expands the existing collection of microbial genomes with paired isolates but also provides evidence of their probiotic and cardiometabolic effects on hosts. This contributes to the mechanistic understanding of host-microbe interactions and paves the way for the development of microbiome-based personalized therapies.

## ACKNOWLEDGEMENTS

We thank the participants and staff of the CGMR for their collaboration and support.

## AUTHOR CONTRIBUTIONS

QZ, LC, WC and XK contributed to conceptualization and funding. The CGMR Consortium Initiative staffs contributed to data and sample collection. LC, QZ, QD and PH contributed to data analysis. QZ, LC, PH and QD contributed to wet lab experiments. PH, QD and YW drafted the manuscript. QZ, LC, YT, SW, CZ, LY, FT, XG, HG, SY, ML, YL, QZ, WL, GW, BY, SC, DH, XW, YJ, LL, QD, BM, TW, HZ, JS, HZ, DF, YS, JZ, LT, HZ, WS, WC and XK contributed to discussion of the content. All authors read and approved the final manuscript.

## LEGENDS OF SUPPLEMENTAL FIGURES

**Figure S1.**
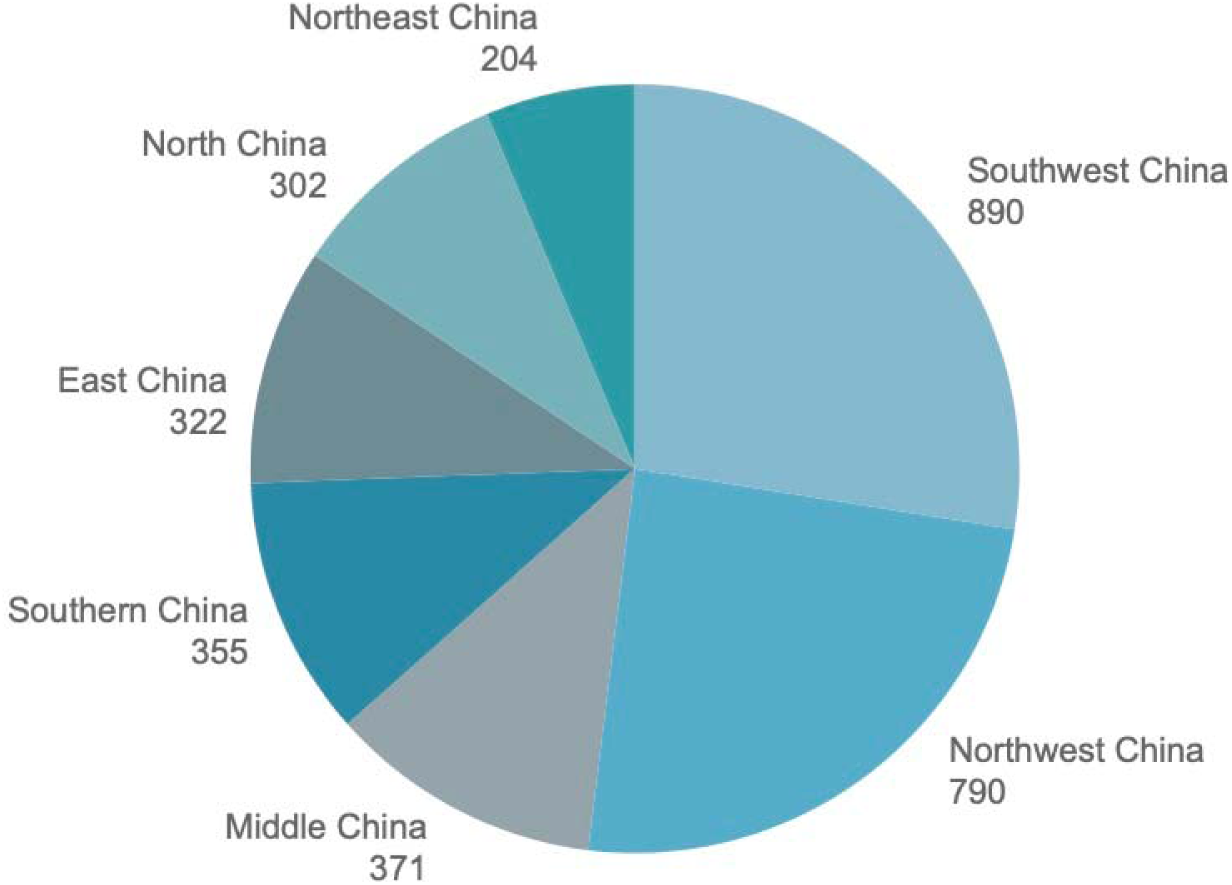
Summary of participant numbers based on 7 geographic regions.

**Figure S2.**
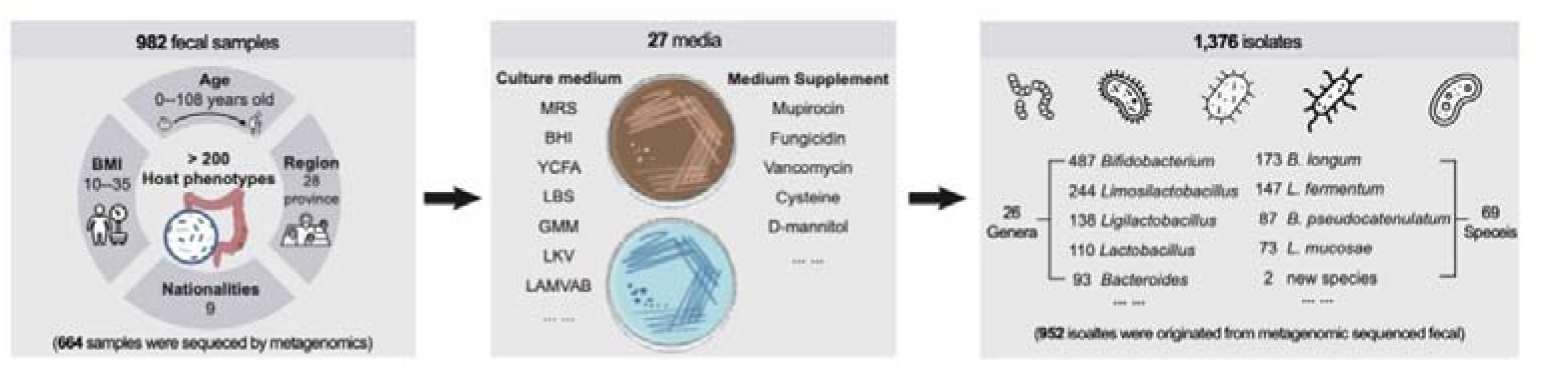
Workflow of bacterial isolation.

**Figure S3.**
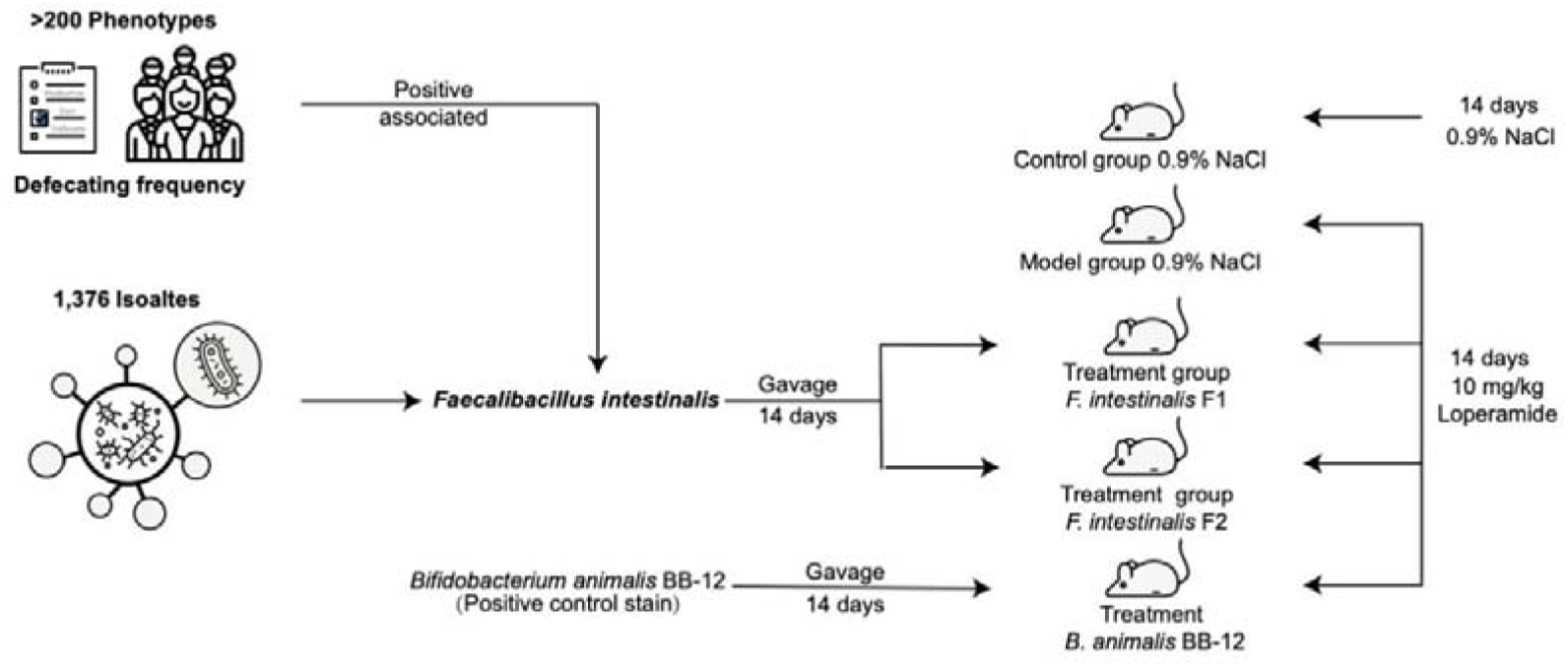
Illustration of the constipation experiment.

**Figure S4.**
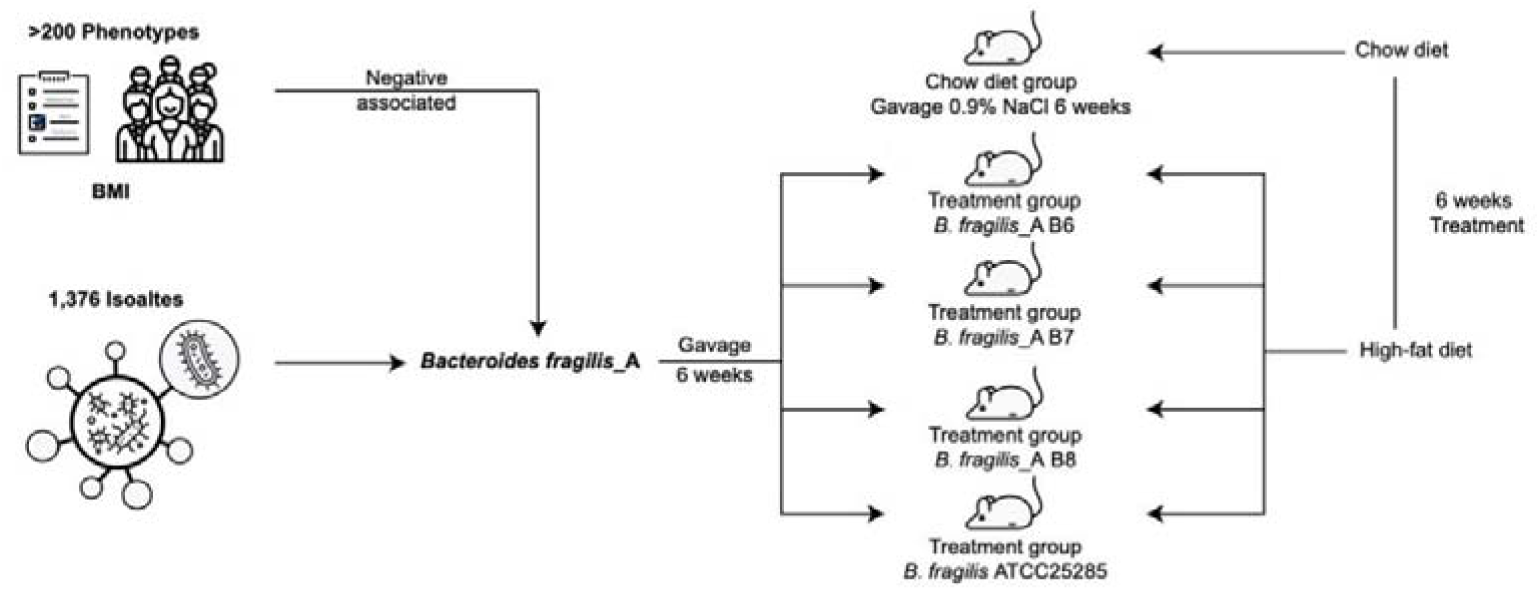
Illustration of the obesity experiment.

**Figure S5.**
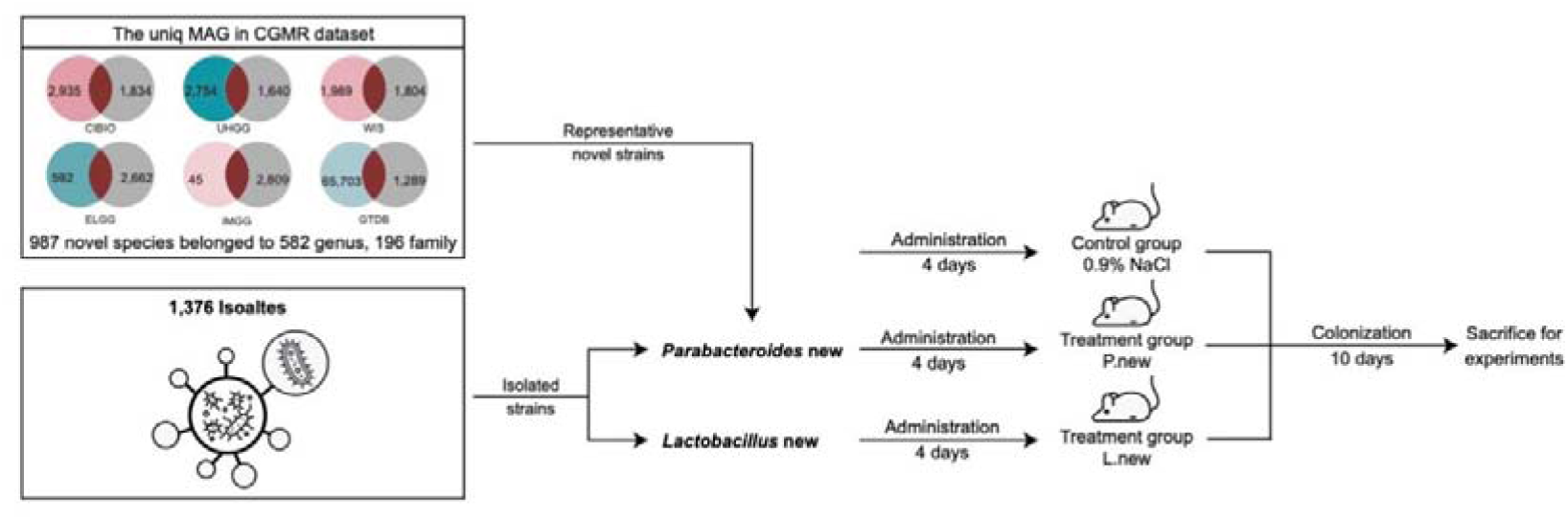
Illustration of the germ-free mice experiment.

**Figure S6.**
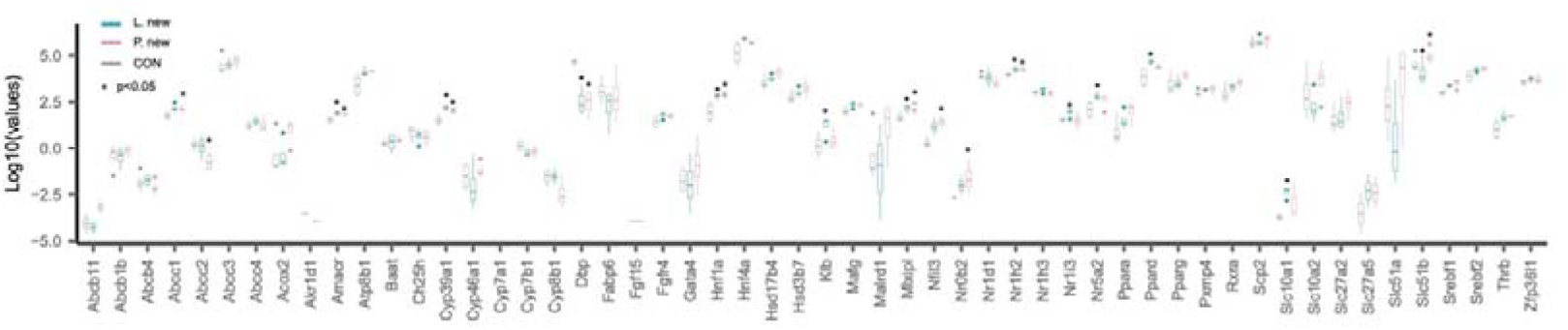
Relative abundance of potential bile acid genes in the colon tissue between groups.

## LEGENDS OG SUPPLEMENTAL TABLES

**Table 1:** Completeness and contamination statistics for 292,550 bins.

**Table 2:** Species-level bins for the CGMR superset dereplicated at 95% ANI.

**Table 3:** Comparison between the CGMR and other public datasets.

**Table 4:** Description of species and annotations with occurrence rate >10% in each of the seven regions in China.

**Table 5:** Co-occurrence network of species-level MAGs.

**Table 6:** Summary statistics of phenotypes.

**Table 7:** Correlation between the existence of MAGs and phenotypes.

**Table 8:** Completeness and contamination statistics for 1,376 isolates.

**Table 9:** MASH distance between isolates and MAGs.

**Table 10:** Functional annotation of 1,376 isolates.

**Table 11:** Summary statistics of phenotypes in obese mice.

**Table 12:** 428 different metabolites in colonic contents between groups.

**Table 13:** 58 out of 258 differential metabolites measured in fermentation *in vitro*.

**Table 14:** Differential serum metabolites of germ-free mice treated with isolates from *Parabacteroides* and *Lactobacillus* species.

**Table 15:** Differentially expressed genes in germ-free mice treated with isolates from *Parabacteroides* and *Lactobacillus* species.

**Table 16:** Gene functional enrichment analysis of differentially expressed genes in *Parabacteroides* new-treated and *Lactobacillus* new-treated groups of germ-free mice.

**Table 17:** Bacterial strains and culture conditions.

**Table 18:** Ingredients of the control diet and high-fat diet fed to the mice.

## STAR *METHODS

### KEY RESOURCES TABLE

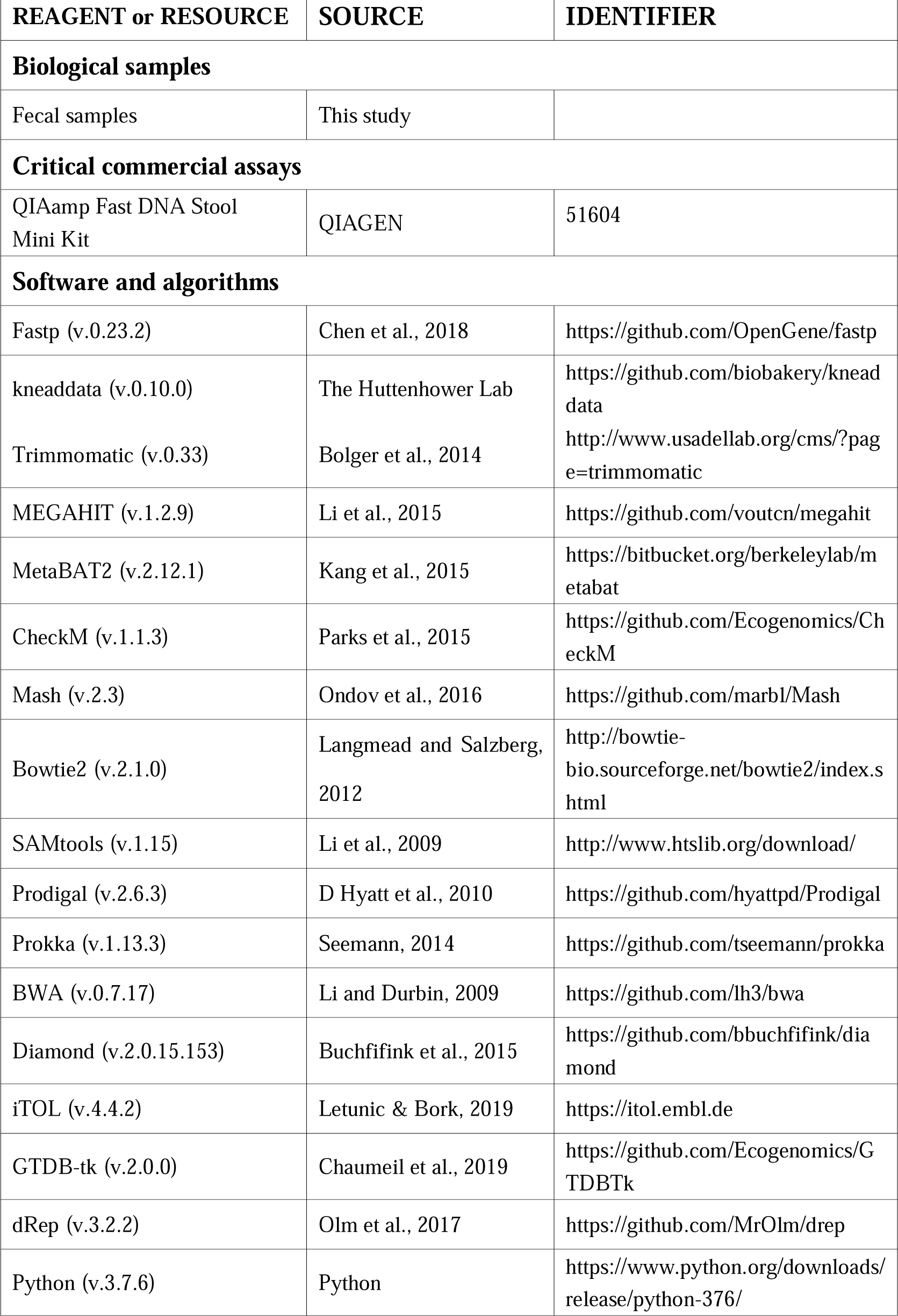
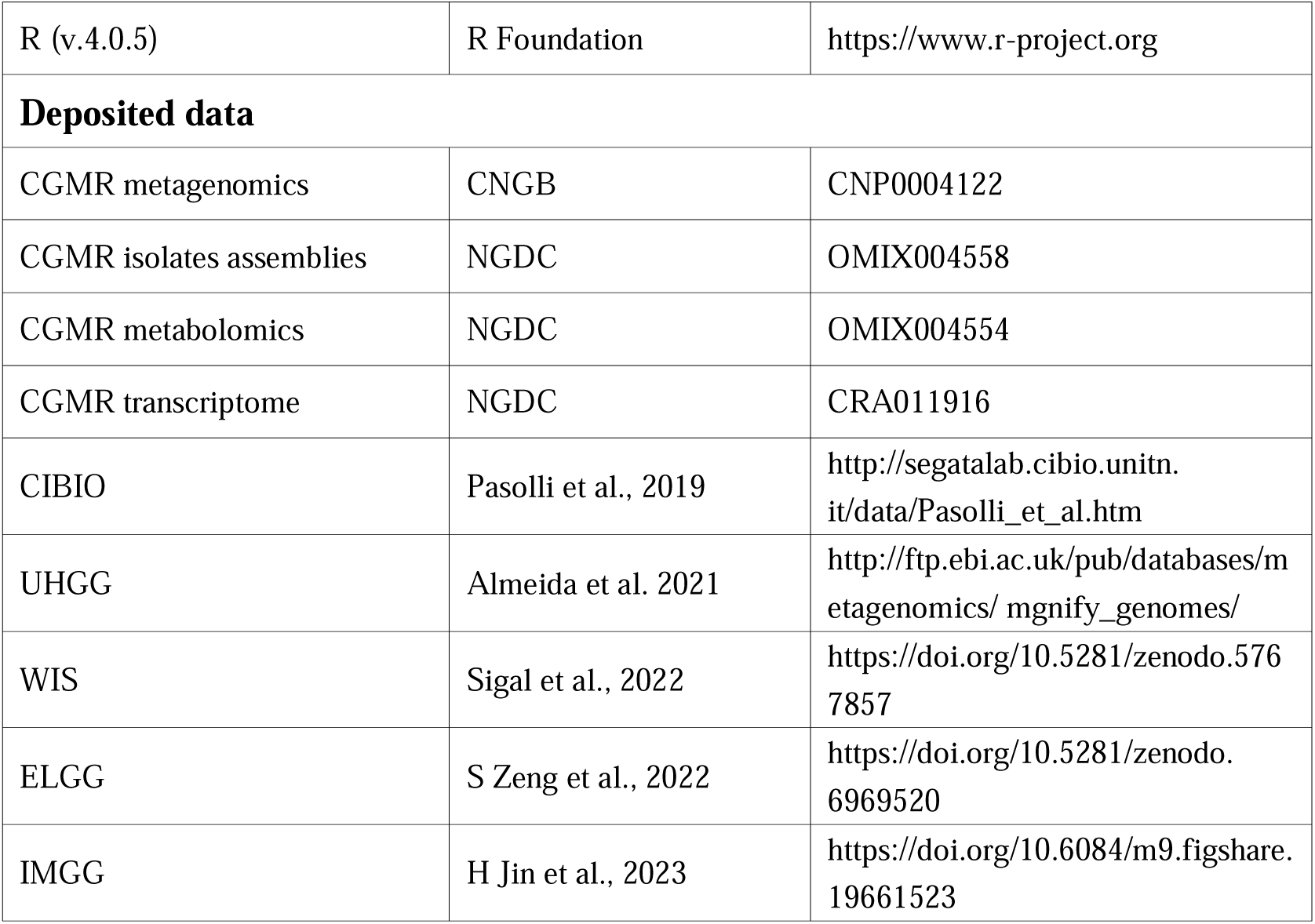

### RESOURCE AVAILABILITY

#### Lead contact

Further information and requests for resources and reagents should be directed to the Lead Contact Lianmin Chen.

#### Data and scode availability

The metagenomic sequencing data used for the analysis presented in this study are available from the China National Gene Bank Data Base (CNGB) under accession id CNP0004122. The genomic assemblies of isolates, metabolome and transcriptome data are deposited in the National Genomics Data Center (NGDC) with project id PRJCA017330. Code used for the analyses is available via https://github.com/MicrobiomeCardioMetaLab/CGMR.database_project.

## EXPERIMENTAL MODEL AND SUBJECT DETAILS

### Human subjects

The study included 3,234 participants from 30 provinces in mainland China (**Figure 1A**). Prior to sample collection, all participants provided informed consent. The study was approved by the ethics review board of Jiangnan University (reference number JNU20220901IRB01). Additionally, a range of phenotypic data were collected from the participants, including anthropometric traits (e.g., age, sex, and body mass index [BMI]), geographical characteristics, residential location (urban and rural), dietary habits, lifestyles, as well as medical history and medication use (**Table S6**).

## METHOD DETAILS

### Metagenomic sequencing

Fecal samples were collected and immediately placed in dry ice within 15 minutes of production. The samples were then transferred to the laboratory. Aliquots of the samples were prepared using a 25% pre-reduced (anaerobic) glycerol solution and 1% cysteine and stored at −80°C until further processing. DNA extraction from the fecal samples was performed using the QIAamp Fast DNA Stool Mini Kit (Qiagen, cat.51604). Following DNA extraction, library preparation and whole-genome shotgun sequencing were conducted using the MGI DNBSEQ-T7 platform for 3,082 samples and the Illumina NovaSeq 6000 platform for 152 samples. During the data processing step, low-quality reads were discarded, and reads originating from the human genome were eliminated by aligning the data to the human reference genome (version NCBI37) using kneadData (v.0.10.0) and Bowtie2 (v.2.1.0) tools ^48, 49^.

### Metagenomic *de novo* assembly and binning

For each of the 3,234 samples, we obtained 5 to 200 million paired-end clean reads with average length of 150 bp. A single-sample assembly strategy was employed to process the data. Initially, the clean reads were used as input for MEGAHIT (v.1.2.9) to perform de novo assembly. This resulted in a total of 5.1×10^8^ contigs longer than 200 bp. Subsequently, Bowtie2 (v.2.1.0) was employed to align the reads back to the filtered assembly, and the resulting alignments were converted to BAM format using SAMtools (v.1.15). To calculate the coverage, we utilized the ‘jgi_summarize_bam_contig_depths’ script from MetaBAT2 (v.2.12.1) on the generated BAM files. These outputs, including the contigs and coverage information, were then utilized for contig binning using MetaBAT2. Contigs shorter than 1.5 kb were excluded during the binning process.

### Genome quality assessing

After the contig binning process, we assessed the quality of the assembled genomes. To do this, we utilized CheckM (v.1.1.3), which relies on the copy-number of lineage-specific single-copy genes. We employed the ‘lineage_wf’ workflow in CheckM to estimate the completeness and contamination of the genomes. We set specific criteria to select genomes that met our quality standards, which included a minimum completeness of 50% and a contamination rate of less than 5% (QS>50, where QS is the completeness-5×contamination). Following this filtering step, we obtained a total of 101,060 medium-quality metagenomic assembled genomes (MAGs) that met our quality criteria.

### Selection of species-level representative MAGs

To identify species-level representative MAGs from the total set of 101,060 genomes, we employed dRep (v.3.2.2) clustering. This process allowed us to extract MAGs that exhibited the highest quality and represented individual metagenomic species. We ran dRep with the following options: ‘-pa’ 0.9 (primary cluster at 90%), ‘-sa’ 0.95 (secondary cluster at 95%), ‘-cm’ larger (coverage method: larger), and ‘-con’ 10 (contamination threshold of 10%). The choice of 95% as the threshold is a common practice for species-level clustering. The genomes were scored based on their completeness, contamination, genome size, and contig N50. From each secondary cluster, only the MAG with the highest score was retained as the winning genome in the dereplicated set. As a result, we obtained a total of 3,707 representative species-level MAGs, which serve as valuable representatives for the metagenomic species in our dataset.

### Taxonomic classification and phylogenetic analyses

Each reconstructed genome was taxonomically classified using GTDB-Tk (v.2.0.0) with the database release 207. The ‘classify_wf’ function was utilized with specific parameters. GTDB-Tk utilizes a set of 120 ubiquitous single-copy proteins to propose bacterial taxonomy. The resulting sequence alignments were then used to generate a maximum-likelihood tree. To visualize and annotate the tree, Interactive Tree Of Life (iTOL, v.4.4.2) was employed.

### Functional annotation of MAGs

For functional annotation, protein-coding sequences (CDS) of each of the 101,060 MAGs were predicted and annotated using Prokka (v1.13.3). Prodigal (v2.6.3) was used within Prokka, applying the following options: ‘-c’ to predict proteins with closed ends only, ‘-m’ to prevent genes from being built across stretches of sequence marked as Ns, and ‘-p single’ to indicate single mode for genome assemblies containing a single species.

### Keystone species-level MAGs

We aimed to identify keystone species that play a crucial role in the organization and stability of the gut ecosystem. These species hold a central position in microbial co-occurrence networks ^1, 23^. To construct the microbial co-occurrence network, we employed Pearson’s chi-squared test based on the presence or absence of species-level MAGs in each sample. In the network, microbial species were considered to co-occur if the number of co-occurrence pairs exceeded the number of co-exclusion pairs. Conversely, if the number of co-occurrence pairs was lower than the number of co-exclusion pairs, the microbial factors were considered to co-exclude each other. To determine significance at FDR< 0.05, we performed 100 permutations. In each permutation, the presence and absence of each species-level MAG were randomly shuffled across the samples. We defined MAGs with high connectivity, representing the top 10% of the network, as keystone species. These keystone species are expected to have a significant impact on the overall structure and functioning of the gut microbiome.

### MAG associations to host phenotypes

The presence or absence of each specie-level MAG was correlated with all the collected phenotypes (**Table S6**). To assess MAG associations to host phenotypes, the Spearman test was applied, and P values were adjusted using the Benjamini-Hochberg (BH) method We examined the correlation between the presence or absence of each species-level MAG and all the collected phenotypes (**Table S6**). To assess the associations between MAGs and host phenotypes, we utilized the Spearman test, which measures the strength and direction of monotonic relationships. The resulting P-values were adjusted for multiple testing using the Benjamini-Hochberg (BH) method ^50^.

### In vitro cultivation of live bacterial isolates

We randomly selected fecal samples from 982 participants and performed bacterial isolation using 27 different culture mediums (**Table S17**). This isolation process allowed us to obtain a total of 1,376 live bacterial isolates. For each isolate, we generated draft genomes for further analysis and investigation.

### Animals

To evaluate the effects of *F. intestinalis* isolates on loperamide-induced constipation, a total of 40 male SPF C57/6J mice (aged six weeks, weighing 19-21g) were included in the study. The mice were housed in controlled conditions with a temperature of 25±2°C, humidity of 55%±5%, and a 12-hour light/dark cycle. The mice were given a one-week adaptation period. The mice were randomly divided into five groups: a control group receiving 0.9% NaCl saline, a Loperamide (LOP) group receiving a gavage dose of 10 mg/kg loperamide to induce constipation once per day for 14 days, and three bacterial intervention groups receiving a gavage dose of 10 mg/kg loperamide along with 1.0 × 10^9^ CFU *F. intestinalis* and *B. animalis ssp. lactis* BB-12 isolates once per day for 14 days. The study protocol was approved by the ethics committee of Jiangnan University (No.20221215c0600125-513).

To evaluate the effects of *B. fragilis_A* isolates on cardiometabolic traits in obesity, a total of 48 male SPF C57/6J mice (aged six weeks, weighing 19-21g) were included in the study. The mice were housed in controlled conditions with a temperature of 25±2°C, humidity of 55%±5%, and a 12-hour light/dark cycle. The mice underwent a one-week adaptation period. The mice were randomly divided into six groups: a control group receiving a normal diet, a high-fat diet (HFD) group (60% fat), and four bacterial intervention groups receiving a 60% fat diet along with a daily gavage of 1.0 × 10^9^ CFU *B. fragilis* FSHXHZ1E1 (B6), FSXLVLS8E1 (B7), FSXLVWS8E1 (B8), and ATCC25285 (BATCC) for six weeks. The detailed diet components can be found in **Table S18**. The study protocol was approved by the ethics committee of Jiangnan University (No.20221130c0700220-471).

To evaluate the effects of two isolates from new *Parabacteroides* and *Lactobacillus* species, a total of 12 male germ-free C57BL/6 mice (aged six weeks, weighing 19-21g) were included in the study. The mice were housed in germ-free flexible film isolators with a temperature of 25±2°C, humidity of 55%±5%, and a 12-hour light/dark cycle. Sterility tests, including culture and PCR, were performed weekly to ensure germ-free conditions. The mice were provided with sterilized food and water irradiated at a high dose of 100 kGy. The two isolates were orally gavaged to the germ-free mice at 8 weeks of age for a duration of 10 days. The study protocol was approved by the ethics committee of Jiangnan University (No.20230228c0300415-039).

All isolates used in the animal experiments were confirmed by 16S sequencing using the 27F and 1492R primers.

### Whole gut transit time in constipation

At the end of each animal experiment (on day 14), the mice were subjected to an overnight fasting period while having access to water ad libitum. The treatment group received lorazepam at a dosage of 10 mg/kg body weight, while the control group received a 0.9% saline solution. After one hour, the mice were administered 250 μL of a 0.5% activated charcoal solution mixed with gum powder to minimize potential irritation to the intestinal tract caused by the activated charcoal. Subsequently, the mice were promptly transferred to a clean and empty separate cage, where they were allowed to eat and drink freely. The whole gut transit time, which represents the time required for the first fecal pellet containing the dye to be expelled and indicates when the feces turn black, was recorded.

### Small intestinal transit time in constipation

Following a 12-hour fasting period, mice in each group were orally administered an activated charcoal solution. Subsequently, the entire small intestine, extending from the pylorus to the cecum, was gently extracted from the abdomen of each mouse and placed on blotting paper. The distance traveled by the activated charcoal along the small intestine was measured, as was the total length of the small intestine. The small intestinal transit was then calculated by determining the percentage of the distance covered by the activated charcoal in relation to the total length of the small intestine for each individual mouse.

### Fecal output in constipation

To assess fecal output over a 5-hour period, the total number of fecal pellets produced by each mouse was recorded for subsequent analysis and comparison. Fresh fecal pellets were individually collected from each mouse using sterile EP tubes. The wet weight of each sample was measured, after which the samples were freeze-dried for 48 hours to obtain the dry weight. The fecal water content of each sample was determined by calculating the difference between the dry weight and the initial wet weight of a representative fecal pellet.

### Measurement of gastrointestinal hormones related to constipation

To quantify the concentrations of motilin (MTL), somatostatin (SS), and vasoactive intestinal peptide (VIP) in mouse serum, ELISA kits from Shanghai Enzyme-linked Biotechnology Co., Ltd. (Shanghai, China) were used following the manufacturer’s instructions. In brief, specific hormone antibodies were coated onto each well of the ELISA plate and incubated with mouse serum at 37°C for 30 minutes. After incubation, the plates were treated with HRP-Conjugate reagent for an additional 30 minutes at 37°C. The reaction was stopped using sulfuric acid, and the absorbance of the reaction mixture was measured at 450 nm using a microplate reader (Multiskan GO, Thermo Fisher Scientific Oy Ratastie 2, FI-01620 Vantaa, Finland) within 15 minutes.

### Determination of short chain fatty acids in constipation

To extract short-chain fatty acids (SCFAs) from fecal samples, 0.05 g of each sample was resuspended in 500 μL of saturated NaCl solution. The mixture was acidified by adding 20 μL of 10% sulfuric acid. Next, 800 μL of diethyl ether was added, and the sample was shaken for 2 minutes. The mixture was then centrifuged at 14,000g for 15 minutes. To remove internal water, 0.25 g of anhydrous Na2SO4 was added, and the sample was stored at −20°C for 30 minutes. Afterward, the supernatant was transferred to a fresh glass vial for analysis using gas chromatography-mass spectrometry (GC-MS). A GC-MS system equipped with a Rtx-Wax column was used for analysis. A 1-μL aliquot of the sample was injected in split mode (10:1). Helium was used as the carrier gas, with a front inlet purge flowrate of 2 mL/min and a gas flow rate through the column of 1 mL/min. The temperature program was as follows: the initial temperature was set at 100LJ for 1 minute, then increased to 140LJ at a rate of 7.5LJ/min, followed by a further increase of 60LJ/min to 200LJ. The temperature was held at 200LJ for 3 minutes. The injection, transfer line, and ion source temperatures were maintained at 240°C, 240°C, and 220°C, respectively. SCFAs, including acetic, butyric, valeric, propionic, and isobutyric acids, were detected using the single scan mode with specific m/z values for each SCFA. To determine the concentration of SCFAs, the response factors for each SCFA relative to 4-methyl-valeric acid were calculated using injections of pure standards. The concentration of SCFAs in the samples was expressed in μmol/g of dry sample.

### Measurement of metabolic traits in obese mice

Throughout the experiment, the body weight of the animals was monitored on a weekly basis. On Day 42, an oral glucose tolerance test was conducted. Prior to the test, the animals were fasted overnight from 6 PM to 6 AM. Then, they received an oral glucose gavage at a dose of 2 mg glucose per gram of body weight. Blood samples were collected at 0, 15, 30, 60, and 120 minutes after the glucose administration. The whole blood samples were centrifuged at 1500 g for 15 minutes at 4°C to obtain the plasma. The upper plasma layer was transferred to new tubes and sent for analysis using an automatic blood biochemical analyzer. The concentrations of glucose, high-density lipoprotein cholesterol (HDL-C), low-density lipoprotein cholesterol (LDL-C), total cholesterol (TC), triglyceride (TG), alanine aminotransferase (ALT), aspartate aminotransferase (AST), alkaline phosphatase (ALP), cholinesterase (CHE), and lactate dehydrogenase (LDH) were determined. On Day 43, the weights of the liver, epididymal fat, and brown fat were measured. Serum leptin and glucagon-like peptide-1 (GLP-1) levels were assessed using a commercial Morinaga Mouse/Rat Leptin ELISA Kit. A 96-well microplate was used for the assay, and the absorbance of the reaction mixture was measured at 450 nm using a spectrophotometer (Multiskan GO, Thermo Fisher Scientific Oy Ratastie 2, FI-01620 Vantaa, Finland) after adding the stop solution, within 15 minutes.

### *In vitro* fermentation of *B. fragilis*_A isolates

Four *B. fragilis*_A isolates (B6, B7, B8, and ATCC25285, 1×10^8^ CFU) were inoculated into anaerobic GMM medium and cultured in 250-ml bottles containing 150 ml of medium. The cultures were incubated at 37°C in anaerobic environment. After reaching the logarithmic growth phase, the bacterial cultures were harvested. To collect the bacterial fluid, the cultures were centrifuged at 12,000 g for 5 minutes. The resulting supernatant, which contains the secreted components, was carefully separated and stored at −80°C for further analysis.

### Metabolome analysis

We have colon feces from obese mice, fasting serum from germ-free mice and *in vitro* fermentation products from *B. fragilis*_A isolates for metabolome analysis. For serum and culture supernatants, 20 μL samples were mixed with 180 μL of 80% methanol and vortexed for 15 seconds. For colon feces, 20 mg samples were mixed with 500 μL of 80% methanol and vortexed three times for 1 minute. All mixtures were then incubated at 4°C for 1 hour to precipitate proteins and subsequently centrifuged at 14,000 g for 15 minutes. The resulting supernatants were collected and stored at - 80°C for metabolome analysis.

The untargeted metabolome analysis was performed using an LC-MS/MS system comprising a ThermoFisher Vanquish UHPLC system coupled with an Orbitrap Q ExactiveTMHF mass spectrometer. The raw data files generated by UHPLC-MS/MS were processed using Compound Discoverer 3.1 (CD3.1, ThermoFisher) software for peak alignment, peak picking, and quantitation of each metabolite. The normalized data was used to predict the molecular formula based on additive ions, molecular ion peaks, and fragment ions. The identified peaks were matched with databases such as mzCloud, mzVault, and MassList to obtain accurate qualitative and relative quantitative results. Metabolite annotations were performed using databases such as KEGG, HMDB, and LIPIDMaps.

To identify differential metabolites between groups, the Wilcoxon test was applied, and the resulting p-values were adjusted using the BH method for multiple testing correction.

### Transcriptome analysis

Colon samples from germ-free mice were collected (0.5 cm distal colon) and the samples were homogenized in TRIzol to extract the RNA. Subsequently, the RNA was converted into complementary DNA (cDNA), and sequencing was performed using the Illumina NovaSeq 6000 platform. The raw paired end reads underwent trimming to remove any low-quality bases or sequencing artifacts, resulting in a set of clean reads. These clean reads were then aligned to a reference genome using the HISAT2 software in the orientation mode to ensure accurate mapping. After alignment, the mapped reads from each sample were assembled using StringTie, which employed a reference-based approach to construct transcripts. The abundance of each transcript was quantified using the transcripts per million reads (TPM) method, which normalizes the expression levels based on the total number of reads and the length of each transcript. Wilcoxon test was applied to compare the expression levels between different groups. Furthermore, functional enrichment analysis of the differentially expressed genes was performed using g:Profiler ^51^.

### Protein structure comparison

The protein structures of potential bile acid metabolism genes identified in isolates from two new species were modeled using AlphaFold2 ^52^. Specifically, the gene *baiA* (EC number: 1.1.1.392) from *Parabacteroides* FGD16K12, and genes *cbh_1* (EC number: 3.5.1.24) and *cbh_2* (EC number: 3.5.1.24) from *Lactobacillus* FHNXY56M7 were subjected to protein structure prediction using AlphaFold2. The resulting protein structures were then compared with confirmed bile acid metabolism proteins using PyMOL. The root-mean-square deviation (RMSD) value was calculated to evaluate the structural similarity between the modeled proteins and the confirmed proteins.

