## Supplementary figures and images for "Gut microbial genomes with paired isolates from China signify probiotic and cardiometabolic effects"

### Supplemental Figure1

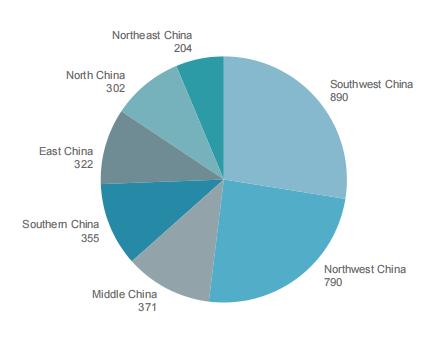

### Supplemental Figure2

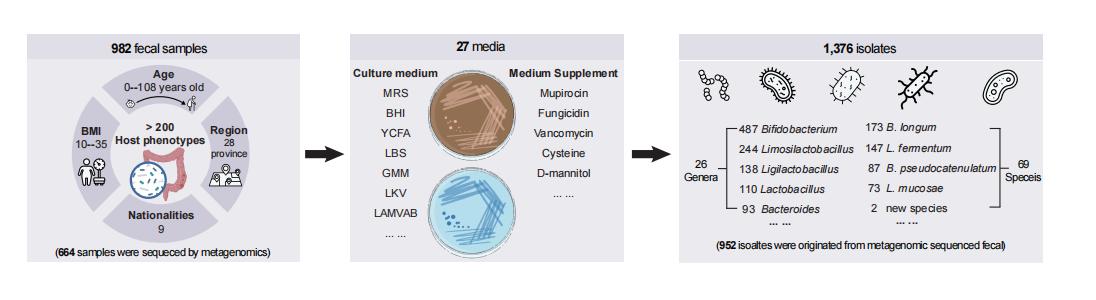

### Supplemental Figure3

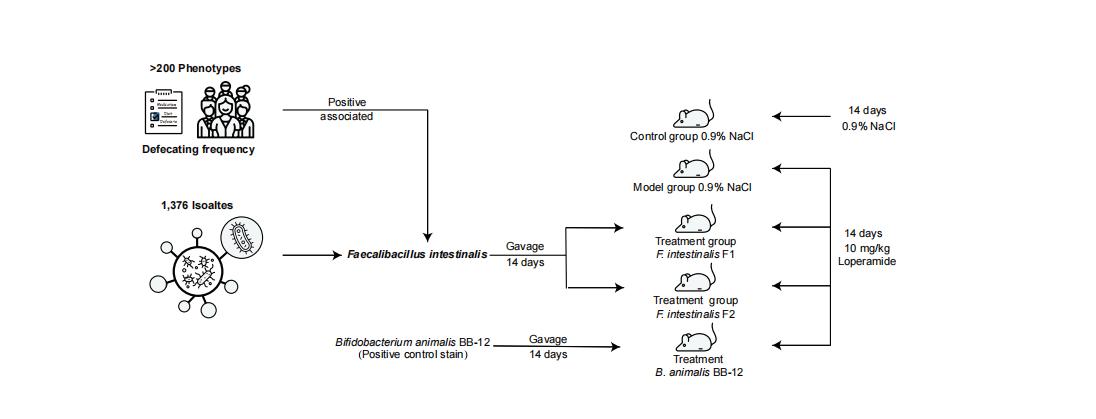

### Supplemental Figure4

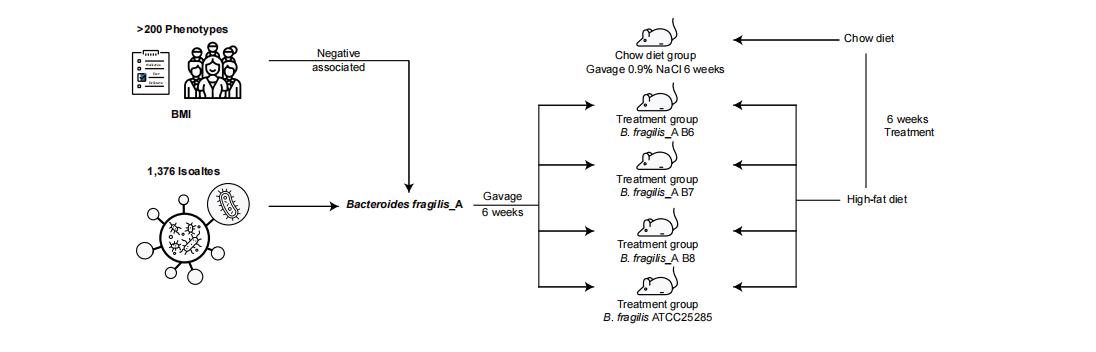

### Supplemental Figure5

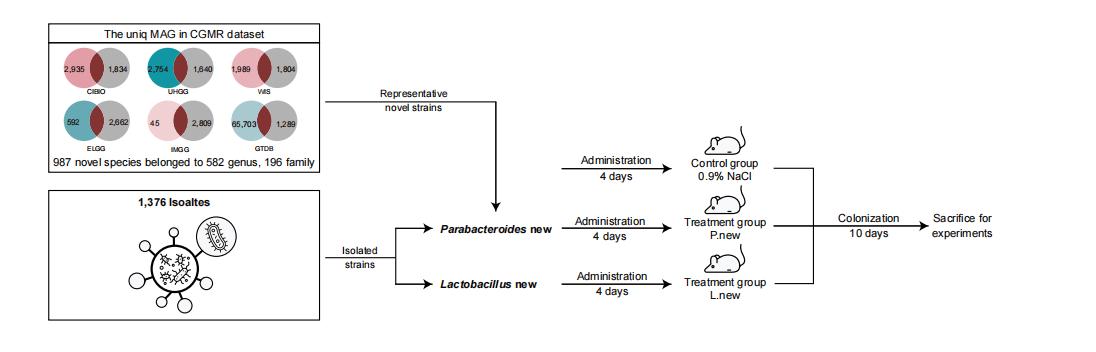

### Supplemental Figure6

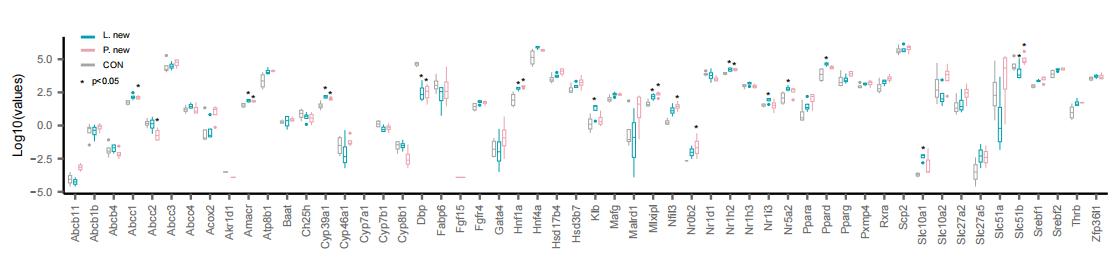
